# Computational Tracking of Cell Origins Using CellSexID from Single-Cell Transcriptomes

**DOI:** 10.1101/2024.12.02.626449

**Authors:** Huilin Tai, Qian Li, Jingtao Wang, Jiahui Tan, Bowen Zhao, Ryann Lang, Basil J. Petrof, Jun Ding

## Abstract

Cell tracking in chimeric models is essential yet challenging, particularly in developmental biology, regenerative medicine, and transplantation research. Existing methods such as fluorescent labeling and genetic barcoding are technically demanding, costly, and often impractical for dynamic or heterogeneous tissues. Here, we introduce CellSexID, a computational framework that leverages sex as a surrogate marker for cell origin inference. Using a machine learning model trained on single-cell transcriptomic data, CellSexID accurately predicts the sex of individual cells, enabling in silico distinction between donor and recipient cells in sex-mismatched settings. The model identifies minimal sex-linked gene sets through ensemble feature selection and has been validated using both public datasets and experimental flow sorting, confirming the biological relevance of predicted populations. We further demonstrate CellSexID’s applicability beyond chimeric models, including organ transplantation and multiplexed sample demultiplexing. As a scalable and cost-effective alternative to physical labeling, CellSexID facilitates precise cell tracking and supports diverse biomedical applications involving mixed cellular origins.

## Introduction

Understanding how a cell’s developmental origin—or cellular ontogeny—influences its function is a central challenge in biology. Accurate tracking of cell origin is essential for studies in developmental biology, regenerative medicine, immunology, and disease modeling [1, 2]. In both experimental and clinical contexts, cell populations of mixed origin frequently arise—for example, in transplantation models, tissue regeneration, or pooled donor samples [3]. Chimeric models, which contain cell populations derived from genetically distinct sources, are a critical tool for such investigations [4]. These models allow researchers to trace cell lineages throughout an organism’s lifespan, offering insights into processes such as development, aging, immune reconstitution, and disease progression. For example, studies of macrophages—immune cells that arise from both prenatal progenitors and postnatal hematopoietic stem cells (HSCs)—have revealed distinct functional roles based on their developmental origin [5]. Prenatal-origin macrophages are seeded into tissues early in development and can self-renew, while HSC-derived macrophages are continuously replenished by blood-derived monocytes originating from the bone marrow [6]. Despite macrophages’ prevalence and functional diversity across all tissues, the specific contributions of these distinct populations of different origins to tissue maintenance, immune responses, and disease remain poorly understood.

Despite its importance, accurate identification of cellular origin at single-cell resolution remains technically challenging. For example, mouse chimeras generated through bone marrow transplantation into recipient hosts of different origins are a frequently employed and powerful tool in biological experimentation. However, existing methods for distinguishing the donor from recipient cells face significant restraints. Traditional approaches rely on tracking genetically engineered artificial markers, such as distinct CD45 alleles or fluorescent reporters. Unfortunately, these markers are often not readily available in the mouse models under study, leading to the need for labor-intensive breeding programs and preparation steps [7, 8]. Additionally, classical methods involving reporter systems designed for lower-throughput analyses often rely on invasive labeling procedures or genetic modifications that are impractical for large-scale single-cell studies or primary human samples. Another strategy which does not require genetic engineering employs sex-mismatched chimeras, using biological sex as a surrogate marker to determine cell origin by detecting either sex chromosomes [9, 10] or their associated gene products [11]. Techniques like fluorescence in situ hybridization (FISH) can detect Y chromosomes in individual cells, providing single-cell resolution, but are only semi-quantitative and incompatible with high-throughput single-cell transcriptomic analysis [9, 10]. On the other hand, PCR-based methods to detect sex-related genes allow for bulk-level quantification of chimerism but lack the single-cell resolution necessary for individual cell origin identification [11]. The inherent limitations of these current techniques hinder our ability to fully characterize and quantify ontogenically distinct cell populations within tissues at the single-cell level, and consequently restrict advances in fields such as immunology, oncology, developmental biology, and regenerative medicine.

To overcome these limitations, we introduce **CellSexID**, an innovative machine learning framework designed to leverage sex as a surrogate marker to identify cellular origin in sex-mismatched chimeric models at single-cell resolution. Unlike prior methods, CellSexID does not require genetic engineering or external labeling and integrates seamlessly into experimental designs by simply including male and female donors or recipients. One of the core functionalities of CellSexID is its ensemble feature selection scheme, which rigorously identifies a minimal yet informative set of sex-linked marker genes that optimize classification performance from a training reference single-cell dataset with male and female cells. By training on our default gene panel, CellSexID achieved robust classification accuracy (AUPRC > 0.94) on a variety of datasets from different tissues and species. CellSexID is also a versatile tool, not limited to chimeric mice model, justified by its accurate prediction in other scenarios like organ transplantation and demultiplexing. Notably, CellSexID generalizes well to our experimentally generated chimeric mouse diaphragm validation dataset, derived from bone marrow transplantation (chimeric mouse diaphragm validation dataset), accurately distinguishing hematopoietic stem cell (HSC)-derived donor (male) from non-HSC-derived host (female) macrophages. Furthermore, we demonstrate that CellSexID’s annotations can enable novel biological insights by applying it to an independent dataset generated from diaphragm macrophages in bone marrow-transplanted mouse (chimeric mouse diaphragm experimental dataset). This analysis revealed that among the 9 macrophage clusters identified in healthy adult skeletal muscle by single-cell RNA sequencing (scRNA-seq) [12], only one was almost exclusively associated with a non-HSC origin, and exhibited a unique anti-inflammatory gene expression profile.

Our findings underscore CellSexID’s ability to uncover ontogeny-specific gene signatures and functional characteristics. This scalable and efficient approach enables high-resolution cell origin tracking across diverse biological contexts. CellSexID holds significant potential for advancing studies of ontogeny-related functions in various cell types and tissues, including applications in chronic inflammatory diseases, cancer, and stem cell therapies, where accurate cell tracking could yield critical insights into disease mechanisms [13]. It is also applicable in diverse settings such as tracking donor and host cells in human organ transplants, distinguishing mixed-sex populations in organoids or pooled samples, and improving sex-aware quality control in large-scale single-cell datasets. Additionally, CellSexID facilitates retrospective sex stratification in public datasets and supports investigations of sex-biased diseases. By enabling precise, single-cell analysis of cellular origin, CellSexID provides a versatile tool for various biomedical research such as developmental biology, immunology, and regenerative medicine, offering new avenues for understanding cellular function, lineage-specific responses, and disease progression in complex tissues.

## Results

### Method overview

CellSexID, as illustrated in Figure 1, first identifies gene features that are predictive of sex using a committee of four machine learning classifiers: XGBoost [14–16], Support Vector Machine [17–19], Random Forest [20–24], and Logistic Regression [25–27]. The committee determines important features based on classifier consensus (a). Using publicly available mouse adrenal gland scRNA-seq datasets containing samples from both sexes [28], the committee identified 14 critical sex-linked markers that enable accurate sex prediction. Based on these selected features, the machine learning classifiers are then trained and predict the sexes for single cells. Validations using public datasets from various tissues and species demonstrated near-perfect prediction performance across all classifiers, highlighting the robustness of the selected features and confirming the feasibility of using single-cell gene expression for sex prediction with machine learning. While all classifiers demonstrated excellent and comparable performance, we selected Random Forest as our default classifier for cell sex prediction. In practice, users can either apply our trained classifier directly or construct a customized model from scratch; both options, along with all associated functionalities, are fully implemented and available in our GitHub repository.

**Figure 1:**
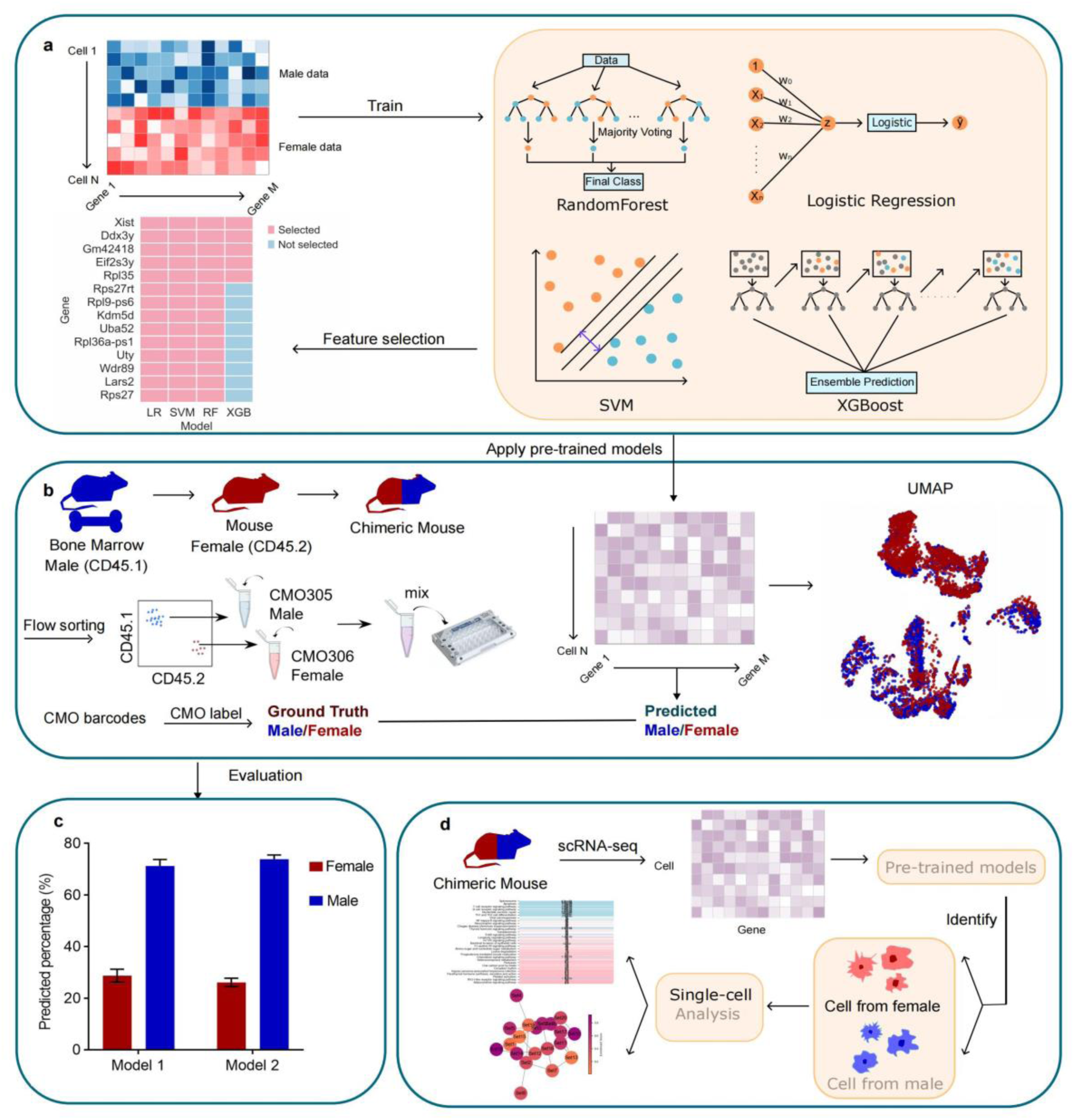
Overview of the CellSexID method. **a,** Identification of sex-specific gene features for predicting cell origin in single-cell RNA sequencing (scRNA-seq) data. A committee of four classifiers—Random Forest, Logistic Regression, Support Vector Machine (SVM), and XGBoost—selects genes important for distinguishing cell sex via majority voting across models. **b,** Experimental validation of CellSexID using chimeric mouse models. Female mice are transplanted with male bone marrow, creating chimeric populations. Diaphragm macrophages are isolated, flow-sorted by CD45 markers, barcoded with cell multiplexing oligos (CMO), and sequenced. The single-cell RNA-seq data from chimeric mice are used to test classifiers trained on a public dataset, with predictions visualized in UMAP space and compared against flow cytometry ground truth for validating CellSexID predictions. **c,** Evaluation of the model’s predictive performance showing predicted cell sex percentages compared to flow cytometry-derived ground truth, providing an assessment of reliability. **d,** Application of CellSexID for annotating cell origin in chimeric mice through a streamlined workflow enables a range of single-cell analyses, supporting studies of cellular dynamics and differences between recipient and donor cells in diverse research contexts.

To evaluate the real-world applicability of CellSexID, we validated the method across multiple public scRNA-seq datasets spanning diverse tissues and species. Additionally, we established a ground-truth system using sex-mismatched chimeric mice generated by bone marrow transplantation, enabling lineage tracing via CD45 allele differences. Macrophages were FACS-sorted, labeled with cell multiplexing oligonucleotides to distinguish male and female origins, and subjected to scRNA-seq following isolation from diaphragm muscle. (Figure 1b). This de novo chimeric mouse diaphragm validation dataset was used to assess the predictive performance of the models trained on the above public datasets in the determination of cell sex origin. Flow cytometry was performed on this dataset to establish ground truth labels, ensuring rigorous evaluation of model predictions. Models trained on the public adrenal gland datasets achieved high accuracy in distinguishing donor (male) from recipient (female) cells when applied to the chimeric mouse diaphragm validation dataset. Standard performance metrics confirmed the generalizability of CellSexID across datasets (Figure 1c).

Finally, to demonstrate its broad utility in biological research, we applied CellSexID to annotate diaphragm macrophages in chimeric mice from a separate mouse chimeric mouse diaphragm experimental dataset (Figure 1d). Differential gene expression (DEG) and pathway enrichment analyses were performed on donor and recipient macrophages, revealing distinct transcriptional profiles and pathways. These findings illustrated the tool’s capacity to accurately track cell origins and facilitate the investigation of transplanted cells in complex biological systems. This controlled chimeric model also highlights how tracking donor and recipient cells provides insights into cell-specific behaviors and disease-associated pathways. Such capabilities can support the development of targeted therapies and enhance our understanding of molecular mechanisms underlying pathogenesis.

### CellSexID identifies a minimal and effective 14-gene marker set for sex prediction through ensemble feature selection

First of all, we justify the necessity of using a machine-learning-based method for selecting multiple gene features and predicting sex by comparing with a rule-based method relying on a single sex chromosome gene. Supplementary Figure 1 shows that our machine learning models significantly outperform a rule-based classifier: *if Uty > 0, then male; otherwise, female*, which is based on the assumption that sex can be determined solely from Y-chromosome gene expression. This is likely due to the challenges posed by high dropout rates and variability in the expression of sex chromosome-linked genes in single-cell RNA-seq data. Such limitations make the genes on sex chromosomes insufficiently reliable for accurate classification without additional features.

For our machine learning classifiers, using a limited number of key gene features for sex-based cell tracking decreases both the complexity and cost of future experimental applications. Therefore, to enhance the real-world applicability of our sex-based cell origin model, we focused on identifying a minimal gene set capable of ensuring accurate sex determination. As shown in Supplementary Figure 2, our feature selection strategy achieves optimal prediction performance compared to models without feature selection or those with varying numbers of top features. Specifically, we compared models in which each committee member selected the top 10, 20, or 200 genes, as well as a baseline using all genes. The final gene panel was derived by selecting genes that at least selected by three committee members. Our results demonstrate that selecting the top 20 genes from each model—resulting in a final panel of 14 genes—achieves the highest predictive performance. Additionally, optimizing feature selection can improve the computational efficiency and accuracy of our model. To identify the most predictive gene features, we employed an ensemble feature selection approach [29–32], combining four machine learning algorithms: XGBoost, Support Vector Machine, Random Forest, and Logistic Regression. Supplementary Figure 3 demonstrates that this ensemble approach outperforms individual models, resulting in superior prediction accuracy.

Our ensemble approach was then trained and tested on a publicly available scRNA-seq datasets from the Gene Expression Omnibus (GEO), featuring mouse adrenal gland immune cell from female (GSM6153751; 3,963 cells) and male (GSM6153750; 2,685 cells) mice [28]. Figure 2a display the top 20 genes with the highest feature importance scores across each model. Figure 2b shows the same results when the feature selection is applied to a human AML BM dataset (GSE289435) [33]. Certain genes consistently ranked as top predictors across different models. By analyzing the committee votes of each gene, we identified a robust set of genes (14 for mouse and 9 for human) selected by at least C = 3 of the 4 models (Figure 2c and d; see Supplementary Figure 4 for justification of this threshold C). These sex-linked gene markers were used in the cell sex predictions in subsequent analyses due to parsimony and their consistently high level performance across multiple algorithms.

**Figure 2:**
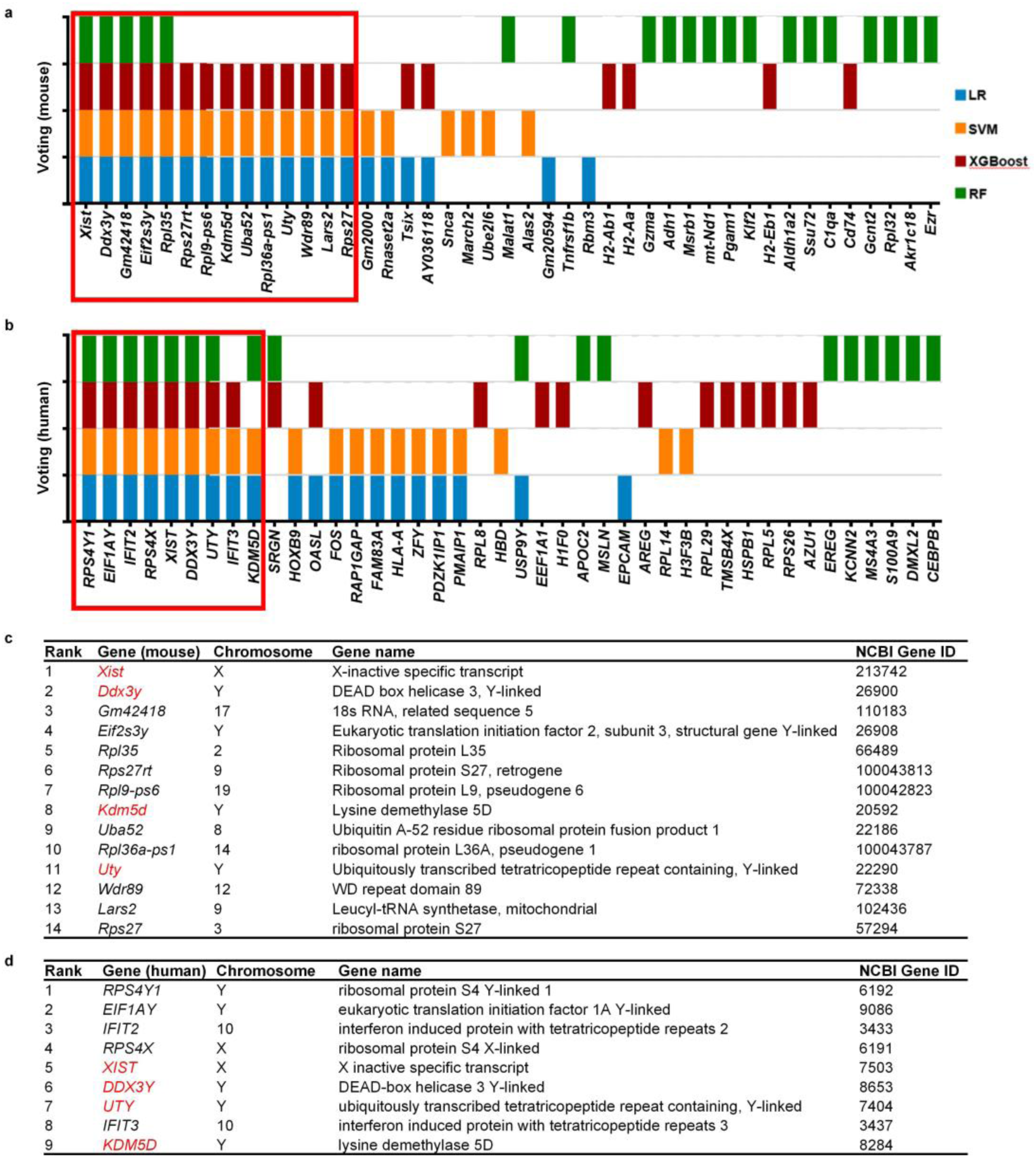
Committee-based machine learning approach identifies key gene markers for sex prediction in mouse and human datasets. **a**, Stacked bar plot showing scaled feature importance contributions from each committee member—Logistic Regression (LR), Support Vector Machine (SVM), XGBoost, and Random Forest (RF)—for mouse gene markers identified using the adrenal gland dataset (GSM6153751 and GSM6153750). The contribution scores are the average of five technical replicates generated using random seeds. In each run, 80% of the data are used for training the model and the rest 20% are for testing and computing the score. Red boxes highlight genes with the highest consensus scores. Feature importance was calculated using min-max normalization within each model, followed by summation across committee members. **b**, Gene markers identified from the human AML bone marrow (BM) dataset (GSE289435, “Single-cell transcriptional atlas of hematopoiesis”). Due to the large dataset size (>80,000 cells), one-fifth of the data was randomly sampled for feature selection in each technical replicates. **c–d**, Final gene marker tables showing top-ranked consensus genes included in the final marker list for mouse (**c**) and human (**d**), including chromosomal locations, full gene names, and NCBI Gene IDs for cross-species ortholog mapping.

It is notable that among the 14 selected genes, only 5 are found on sex chromosomes whereas the rest are autosomal. The sex chromosome-linked genes have a combination of X chromosome (*Xist*) and Y chromosome (*Ddx3y*, *Eif2s3y*, *Uty*, *Kdm5d*) locations. *Xist* plays a central role in X chromosome inactivation in females [34]. *Ddx3y* and *Eif2s3y* dictate male-specific functions such as spermatogenesis [35, 36]. *Uty* [37] and *Kdm5d* [38], located on Y chromosome, are also recognized as reliable sex markers, with their expression limited to males and their roles linked to histone modifications and chromatin organization [39]. Among the 9 autosomal genes identified, several are also known to exhibit sexual dimorphism in other cell types [40, 41]. To further validate our findings, we compared the performance of our 14 selected genes against features limited to only the Y chromosome or the two sex chromosomes together. As shown in Supplementary Figure 5, the genes selected by our model, including those located on autosomal chromosomes, led to a superior predictive performance. Specifically, the “All Chromosomes” condition includes all 14 selected gene features (selected using our feature selection strategy from all genes on all chromosomes), while the “Established sex markers from selected” condition uses only the sex-linked genes from the 14-gene set. This comparison between “All Chromosomes” and “Established sex markers from selected” is exactly the comparison requested by you, and it provides a clear justification for the inclusion of autosomal genes. The results clearly show that models using the full 14-gene set outperform those restricted to sex chromosome-linked genes. In this regard, we applied the same feature selection procedure to sex-linked genes alone “X and Y Chromosomes Only” and Y chromosome genes alone “Y Chromosome Only”, and both subsets exhibited inferior performance compared to the first bar with our 14 features selected from all genes. This underscores the value of our end-to-end feature selection approach, powered by the machine learning model committee, in identifying robust sex markers beyond sex chromosomes. A more comprehensive list of identified features is available in Supplementary Table 1.

In addition, to determine whether these genes have been previously implicated in sex determination or related biological processes, we performed literature searching and found that all the autosomal genes in the sex-linked marker list of human and mouse can find evidence of sex relevant in the literature: For mouse: *Gm42418*[42], *Uba52* [43], *Wdr89* [44], *Lars2* [45], and *Rps27* [46]. For example, *Uba52* exhibits sex-specific regulation in the amygdala during fear memory formation [38], and *Wdr89* shows sex-biased expression in the postnatal pituitary gland, implicating its potential role in hormonal regulation [39]. *Lars2* has been identified as a gene with sex-dimorphic expression across multiple cell types, as revealed by single-cell transcriptomic analyses [40]. For human, *IFIT2* [47, 48] and *IFIT3* [48, 49] are interferon-stimulated genes with higher expression in females and well-established roles in immune responses with known sex differences. In addition, the literature also suggests that it is common for autosomal genes to be relevant to sex. For example, *Nr5a1* (on mouse chromosome 2) is known to play a critical role in sex determination and development [50–52]. With these identified autosomal genes in the sex-linked marker lists, we also performed enrichment analysis using Toppgenes [53]. PubMed gene sets (Enrichr, version 2023), as shown in Supplementary Figure 6. Statistically significant terms show relevance to sex difference, such as male germ cell-specific ribosome and translation-related pathways, which are functionally associated with sex-biased expression and male fertility.

In summary, our ensemble feature selection approach not only identified a biologically relevant and minimal gene set but also demonstrates the power of combining multiple machine learning algorithms for robust and reproducible feature selection. This consensus-based selection method was able to restrict the number of predictors to an easily manageable level without sacrificing performance. In addition, to provide more flexibility, in CellSexID released in our GitHub repository we also provide the functionalities allowing the users to directly provide their own gene list or performing feature selection on their own datasets.

### Validations using datasets from various tissues and species demonstrate CellSexID’s accurate sex prediction based on the robust feature selection strategy

To further assess the effectiveness and generalizability of our feature selection approach, we validated its performance across a diverse range of datasets spanning multiple tissues and species. These validations included (i) testing the default set of 14 sex-linked markers on various datasets, (ii) evaluating the robustness of our feature selection pipeline by applying it to new datasets, and (iii) demonstrating applicability across species using human data.

We first conducted a baseline cross-validation using the publicly available mouse adrenal gland dataset from Dolfi et al. (GSM6153751 for female mice and GSM6153750 for male mice). In this dataset, mouse adrenal gland dataset (CD45⁺ hematopoietic cells) were isolated from adrenal glands, and scRNA-seq was performed using the 10x Genomics platform. The sex origin of the cells was definitively known based on the source animals. Using an 80/20 train-test split, we trained and evaluated our machine learning models on this dataset (Figure 3a). All models demonstrated consistently high and comparable performance across multiple metrics with those 14 selected features. Our default classifier, Random Forest achieved an AUROC of 0.9723, supporting both the validity of our marker selection and the strong baseline performance of CellSexID.

**Figure 3:**
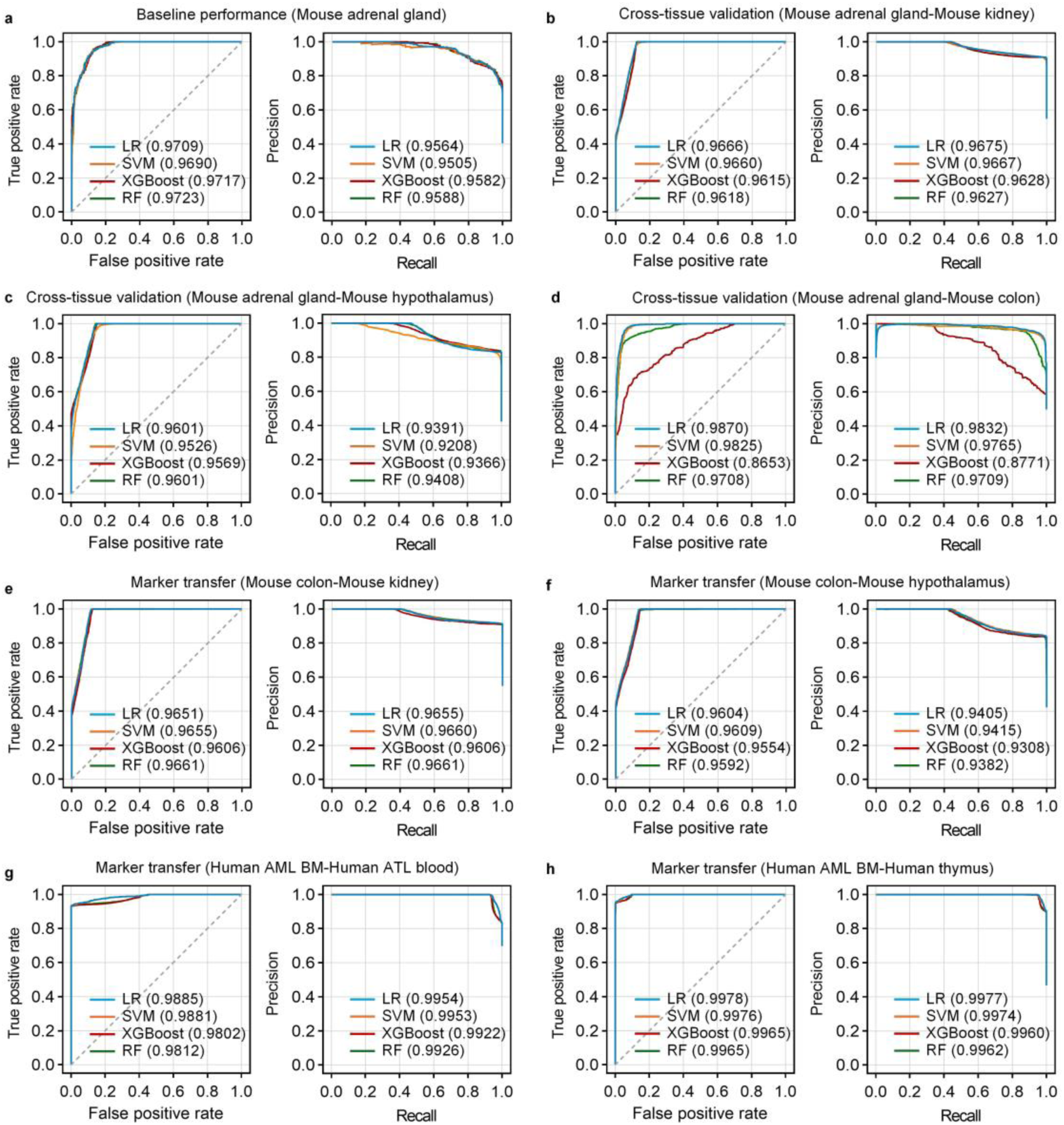
CellSexID performance evaluation across tissues and species. **a,** Baseline performance of four machine learning models—Logistic Regression (LR), Support Vector Machine (SVM), XGBoost, and Random Forest (RF)—trained on the public mouse adrenal gland dataset. Internal validation was performed using an 80/20 train-test split, with 14 sex-linked marker genes selected from the same dataset. **b,** Cross-tissue validation (Kidney): Models trained on the adrenal gland dataset were tested on a mouse kidney dataset (GSE129798), using the adrenal-derived marker list. **c,** Cross-tissue validation (Hypothalamus): Models trained on the adrenal gland dataset were tested on a mouse hypothalamus dataset (GSE201032), using the adrenal-derived marker list. **d,** Cross-tissue validation (Colon): Models trained on the adrenal gland dataset were tested on a mouse colon dataset (GSM2967048), using the adrenal-derived marker list. **e,** Marker transfer in mouse – Kidney test: A new marker list was selected using our feature selection pipeline applied to the mouse colon dataset (GSM2967048). Based on this colon-derived marker list, models were trained on the colon dataset and tested on the kidney dataset. **f,** Marker transfer in mouse – Hypothalamus test: Using the colon-derived marker list, models trained on the colon dataset were tested on the hypothalamus dataset. **g,** Marker transfer in human – T-cell leukemia test: Models were trained on the human acute myeloid leukemia (AML) bone marrow (BM) dataset (GSE289435) using 9 human sex-linked markers and tested on a human adult T-cell leukemia/lymphoma (ATL) blood dataset (GSE294224). **h,** Marker transfer in human – Thymic epithelial test: Models trained on the human AML BM dataset (GSE289435) using the same 9 sex-linked markers were tested on a human thymic mimetic epithelial cell dataset (GSE262749).

Then, to further validate the robustness of our sex-linked markers and the flexibility of our method, we then tested whether the machine learning models trained on our default mouse adrenal gland dataset with the default 14 sex-linked markers can maintain accurate sex prediction on other very different datasets from other mouse tissues. In Figure 3b-d, we present these cross-tissue validation results of training our machine learning classifiers with the selected sex-linked gene markers on the mouse adrenal gland dataset and testing on a mouse kidney (GSE129798) [54], a mouse hypothalamus-PVN (GSE201032) [55], and a mouse colon (GSM2967048) [56] dataset. The consistent high prediction performance in these new settings (most AUROC >0.95) further validate the robustness and the generalizability of CellSexID and our selected sex-linked marker list.

To evaluate cross-tissue generalizability, we trained new models using the mouse colon (GSM2967048) [56] dataset and applied the same methodology to identify 11 markers from this tissue. We then tested the model performance using these markers on scRNA-seq datasets from other mouse tissues: mouse kidney (GSE129798), and mouse hypothalamus (GSE201032). As shown in Figure 3e–f, CellSexID maintained high prediction accuracy across these datasets, with Random Forest AUROC values exceeding 0.96. These results further demonstrate the robustness of our marker set and the transferability of models trained on one tissue to other distinct tissues.

Next, to test generalizability across species, we applied our feature selection and prediction pipeline to human data. Feature selection was performed on a human bone marrow mononuclear cells(BMMCs) from acute myeloid leukemia (AML) dataset (GSE289435), and the resulting markers were used to train models that were then validated on independent human datasets, including human peripheral blood mononuclear cells (PBMCs) from T-cell leukemia–lymphoma (ATL) patients (GSE294224) [57] and human thymus cells (GSE262749) [58] datasets. As shown in Figure 3g–h, the models achieved excellent predictive accuracy across these human datasets, further demonstrating the robustness and generalizability of CellSexID across species.

Together, these results show that (i) our default 14 sex-linked gene markers perform well across tissues, (ii) feature selection is robust to dataset choice, consistently yields strong classifiers, and (iii) CellSexID is broadly applicable across species, tissues, and experimental conditions. These validations collectively demonstrate that CellSexID is a reliable and broadly applicable tool for sex-based cell origin inference, offering a scalable and label-free alternative for diverse single-cell transcriptomic studies.

### Validation of CellSexID for accurate cell origin tracking using cell sorting

After demonstrating the strong sex prediction performance of CellSexID on public datasets, we next evaluated its applicability for tracking cell origin in a real-world setting by performing “de novo validation” in sex-mismatched chimeric mice. The chimeric mice were created by transplanting bone marrow from male donor mice (CD45.1 allele) into female (CD45.2 allele) host recipient mice (Figure 4a). The resident macrophages present in the diaphragm muscle of female recipient mice were protected from the pre-transplant irradiation protocol through the use of lead shielding as we have recently described [59]. At 8 weeks post-transplantation, macrophages were sorted and pooled from the diaphragms of two chimeric mice by FACS based on their CD45 allele status (CD45.1 or CD45.2) and then labelled with Cell Multiplexing Oligo (CMO) barcodes. Using our scRNA-seq analysis pipeline, these distinct CMO barcodes established the ground truth for sex-based macrophage origin, providing a robust benchmark for experimentally evaluating CellSexID’s annotation performance in this “chimeric mouse diaphragm validation dataset”.

**Figure 4:**
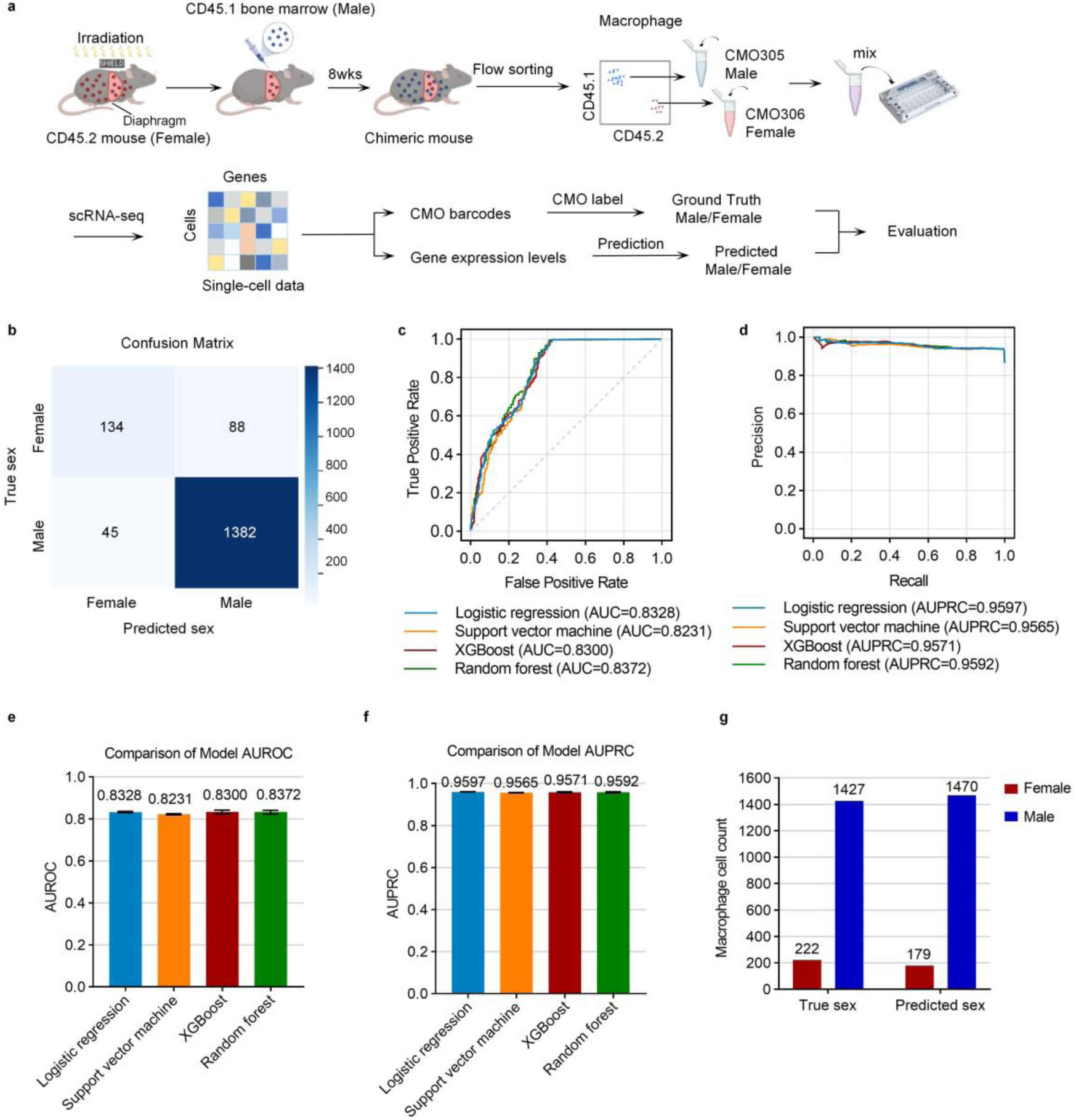
Reliable performance of cell origin classifiers on additional experimentally generated single-cell validation data from a chimeric mouse with cell sorting as ground truth. **a,** Generation of a chimeric mouse diaphragm validation dataset by creating chimeric mouse, collecting single-cell data, and using cell sorting as ground truth. A female CD45.2 mouse, irradiated with diaphragm shielding, was transplanted with male CD45.1 bone marrow. After 8 weeks, diaphragm macrophages were isolated, stained to identify live macrophages (CD11c-, SiglecF-, F4/80+, CD11b+), flow-sorted by CD45.1 and CD45.2 expression, barcoded (CD45.2: CMO306, CD45.1: CMO305), pooled, and sequenced using 10x Genomics. **b**, Confusion matrix of Random Forest predictions based on the 14 sex-linked markers, demonstrating high classification accuracy for cell origin. **c**, AUROC scores of all four classifiers (trained on the public dataset), confirming reliable performance. **d**, Precision–recall (PR) curves for the same classifiers, showing consistent predictive robustness. **e–f**, Comparative analysis of AUROC and AUPRC values for each classifier using bootstrap resampling. Error bars represent 95% confidence intervals (2.5th–97.5th percentile) from 100 bootstrap iterations with replacement applied to the training data. In each iteration, models were retrained on the resampled training set and evaluated on the fixed test set. All models achieved AUROC > 0.82 and AUPRC > 0.95. **g**, Bar plot comparing Random Forest-predicted cell origin ratios (based on sex) with ground truth, illustrating the classifier’s high accuracy using the selected sex-linked markers.

The results of this validation, summarized in Figure 4b-g, demonstrate the high accuracy of CellSexID in classifying cell origin. Figure 4b shows the confusion matrix for the best-performing model, Random Forest, highlighting its precision and recall by detailing true positive, true negative, false positive, and false negative rates. Figure 4c presents the ROC curves for the four machine learning algorithms, illustrating each model’s ability to distinguish between male and female cells, with the area under the curve (AUC) as a primary metric of performance. Figure 4d shows the PRC curves for the same models, offering a balanced perspective on precision and recall by evaluating the trade-off between false positives and false negatives. To further assess model stability, Figures 4e and 4f present bootstrapped AUROC and AUPRC performance across multiple samples for each model. These bar plots confirm the robustness of the scores across all classifiers. Detailed confusion matrices and performance metrics for all four classifiers are provided in Supplementary Figure 7 a-b confirming that the robust performance is not limited to a single algorithm but reflects the effectiveness of our feature selection approach.

Finally, Figure 4g compares the predicted male-to-female ratio with the actual true ratio based on the chimeric mouse diaphragm validation dataset, underscoring the accuracy of CellSexID in estimating population composition in a real-world setting. This comprehensive evaluation of CellSexID with its minimal 14-gene set in the de novo chimeric mouse diaphragm validation dataset supports its reliability for accurate cell origin tracking across multiple scRNA-seq scenarios. Notably, the models were trained on a public mouse adrenal gland dataset and tested on the independent bone marrow transplantation-derived chimeric mouse diaphragm validation dataset, yet still achieved high predictive accuracy. This cross-dataset generalizability serves as strong evidence of CellSexID’s robustness. Importantly, CellSexID operates without relying on assumptions about specific tissue types, cellular contexts, or even species, making it broadly applicable to diverse experimental systems and research questions.

To provide a more comprehensive view of the model’s performance, we calculated class-specific metrics in Supplementary Figure 8. In this supplementary figure, AUROC and AUPRC for male and female predictions to provide a more detailed view of the model performance, showing decent performance in predicting both sexes. To better understand the performance and investigate the characteristics of the misclassified cells, we performed comparative analysis between the correctly classified and misclassified cells. We discovered that misclassified cells tend to be noisier and have statistically significant higher dropout rate than correctly classified cells (Supplementary Figure 9). A higher dropout rate (zero expression) of genes is a kind of typical technical noise in single-cell data, and it naturally leads to the loss of information of the sex-linked gene, and thus might misleads the classification. In this analysis, we compared correctly classified cells versus misclassified cells within female and male groups, focusing on the 14 sex-linked gene markers used by our machine learning classifiers. (Supplementary Figure 10) In female cells, correctly classified cells showed significant upregulation of *Xist*, which is expected and consistent with the expected high expression of *Xist* in female cells due to X chromosome inactivation [60]. In contrast, among female cells that are misclassified as male cells, the loss of expression of *Xist* implies that they are classified as male cells might be due to the key female marker was not detected. In male cells, correctly classified cells showed significant upregulation of *Lars2* and *Uba52*, two genes previously reported to be more highly expressed in male cells [41, 61]. The low expression or dropout of these genes in male cells may lead to insufficient male-specific transcriptional signatures, thereby increasing the chance of being misclassified as female cells. Overall, these observations are consistent with a well-known property of scRNA-seq data: low mRNA capture efficiency often leads to frequent “dropout” zeros, which can affect the accurate classification of cell identity [62].

Taken together, these findings validate the reliability of CellSexID in real-world applications. Its strong performance on the independently generated chimeric mouse diaphragm validation dataset—despite being trained on a separate tissue—demonstrates robust generalization. By accurately identifying cell origin under realistic experimental conditions, CellSexID proves to be a dependable tool supporting biological investigations where traditional labeling is not feasible or cost-effective.

### CellSexID is versatile and capable of cell origin tracking in various application scenarios

While our validations have been focused on chimeric models, CellSexID’s capability of cell origin tracking based on sex prediction enables a wide range of downstream applications without requiring genetic modification, physical barcodes, or antibody-based labeling. To demonstrate this, we explored its performance in two additional experimental contexts—organ transplantation and sample demultiplexing.

We tested CellSexID’s utility in distinguishing donor- and recipient-derived cells in organ transplantation contexts using sex-mismatched models. Figure 5a illustrates results from human organ transplantation experiments, in which our default Random Forest classifier trained on the human AML BM dataset (GSE289435), based on our selected 9 human sex-linked markers, accurately differentiated donor cells from recipient cells in a human kidney transplant dataset (GSE151671) [63] based solely on sex-specific gene expression patterns. This allowed clear and reliable tracking of cellular origins without relying on traditional genetic modifications or invasive labeling methods. The high predictive performance (over 0.9 AUROC) demonstrated in this experiment underscores CellSexID’s robustness and precision in complex in vivo transplantation scenarios. In addition, while the original study identified recipient-derived fibroblasts as key contributors to graft fibrosis, CellSexID enabled a more accurate, complete, and scalable classification of cell origin, reducing ambiguity in sex-mismatched transplant samples and reinforcing this critical biological insight with greater confidence, as justified in Figure 5a, where CellSexID-selected markers lead to higher performance than the ones found in the original study. These results illustrate that CellSexID can effectively address critical biological and clinical challenges in transplantation biology, such as immune reconstitution studies, organ engraftment analyses, and monitoring cell lineage dynamics in vivo. Importantly, this capability extends directly into clinical contexts (including human transplantations), where donor-recipient sex mismatches frequently occur. Thus, these results significantly extend the applicability of CellSexID beyond traditional animal chimeric studies, highlighting its utility as a broadly relevant tool for both basic biomedical research and clinical translation.

**Figure 5:**
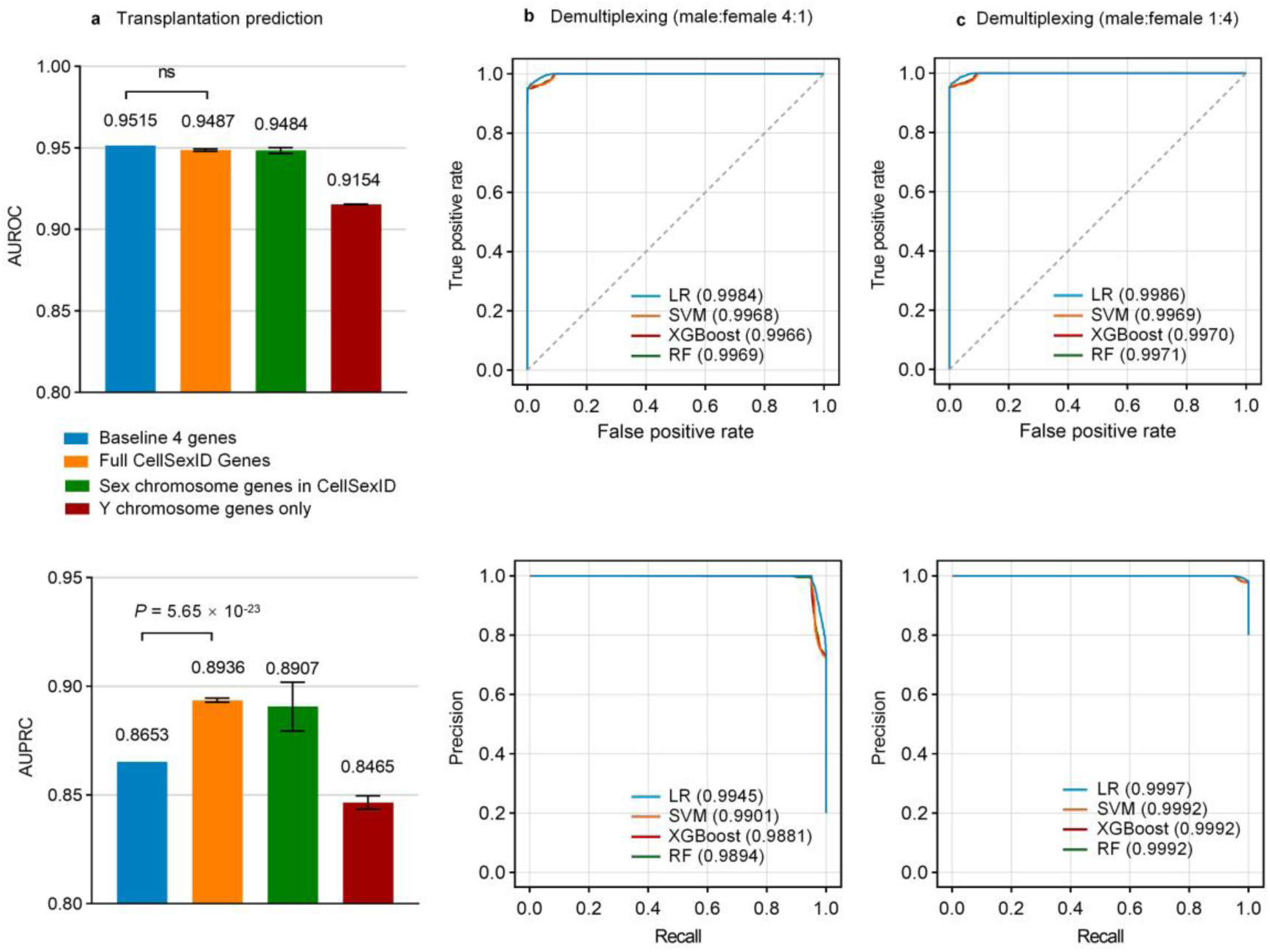
CellSexID enables accurate cell origin tracking in organ transplantation and demultiplexing scenarios. **a,** Performance evaluation for transplantation prediction using AUROC and AUPRC metrics. Random Forest classifiers were trained on human AML bone marrow mononuclear cells (GSE289435) using different gene marker sets and tested on human kidney transplantation data to distinguish donor from recipient cells. Four marker sets were compared: baseline 4 genes (*XIST, RPS4Y, EIF1AY, DDX3Y* highlighted by original dataset providers), full CellSexID gene set, sex chromosome genes within CellSexID, and Y chromosome genes only. P-values from the Two One-Sided Tests (TOST) indicate that the CellSexID marker sets achieve comparable AUROC and significantly superior AUPRC compared to the baseline method, suggesting overall better performance. **b-c,** Demultiplexing performance on synthetic datasets with skewed sex ratios of 4:1 male:female (b) and 1:4 male:female (c), created by sampling 12,000 cells from human thymic medullary epithelial cells dataset (GSE262749). ROC curves (top) and Precision-Recall curves (bottom) demonstrate near-perfect classification performance across all machine learning models (Logistic Regression-LR, SVM, XGBoost, Random Forest-RF) with AUC values >0.98, indicating robust performance even under challenging class imbalance conditions.

To further showcase CellSexID’s general applicability, we demonstrate that, by predicting sex as a surrogate target and using sex-mismatched single-cell multiplexed samples, CellSexID can perform the single-cell demultiplexing task and significantly reduce costs and labor required for traditional multiplexing approaches. CellSexID eliminates the need for physical barcodes or antibody-based panels by leveraging endogenous gene expression for sample demultiplexing. It operates without additional reagents or protocol modifications, significantly reducing experimental complexity, reagent costs, and hands-on time compared to traditional multiplexing approaches such as antibody hashing.

To justify CellSexID’s ability to reliably discriminate cell origins in samples containing mixtures of male and female cells from multiple individuals processed simultaneously, synthetic datasets were created with extreme male:female imbalances, including near 1:4 (2,400 male : 9,600 female cells) and 4:1 (9,600 male: 2,400 female cells) ratios among five different synthetic compositions totaling 12,000 cells each. These synthetic datasets were generated by systematically sampling from the human thymus dataset (GSE262749): Thymic mimetic cells in humans – medullary thymic epithelial cells dataset to create controlled mixtures that simulate hard-to-predict scenarios where male and female cells from different individuals are processed together in a single sequencing run.

The model was trained on the human AML BM dataset (GSE289435) dataset containing 87,171 cells and tested on these synthetic imbalanced datasets using a panel of 9 sex-linked marker genes (*RPS4Y1, EIF1AY, XIST, DDX3Y, UTY, KDM5D, IFIT3, IFIT2, RPS4X*). CellSexID successfully distinguished cells solely based on their sex-linked expression patterns across all synthetic ratios (Figure 5b and c), achieving near-perfect performance with AUROC scores exceeding 0.996 and accuracy rates above 96% even in the most extreme imbalanced scenarios, demonstrating robust demultiplexing capabilities without relying on traditional physical barcodes or antibody-based multiplexing approaches. This approach validates that sex-linked gene expression provides a reliable biological signature for cell origin determination across diverse mixing ratios and population compositions, and is therefore applicable to real-world demultiplexing scenarios.

We further emphasize that our previously described validation experiment (Figure 4)—in which male and female cells from two mice were pooled and sequenced together—serves as a real-world demonstration of barcode-free demultiplexing. In this setting, externally added barcodes (CMOs) provided the ground truth, allowing us to directly test whether CellSexID could accurately assign cell identities based solely on sex-linked gene expression. All four machine learning models achieved strong annotation performance, with AUROC values exceeding 0.82 and AUPRC values above 0.95. These results validate CellSexID’s cell origin tracking performance and demonstrate its practical utility as a cost-effective and robust alternative to traditional barcode-based multiplexing approaches.

Collectively, these examples clearly illustrate that CellSexID is broadly applicable beyond conventional animal chimeric models. Specifically, by leveraging inherent biological differences encoded in sex-linked gene expression, CellSexID can reliably differentiate cell origins, track cell dynamics, and facilitate innovative multiplexing strategies. Compared to conventional methods, CellSexID requires less effort and budget as an in silico approach. Additionally, these tested scenarios directly align with broader applications such as quality control for single-cell datasets, retrospective sex-specific analyses of public datasets, and studies investigating sex-associated mechanisms in human disease. This versatility positions CellSexID as a valuable, cost-effective tool suitable for diverse biological, clinical, and bioinformatic applications.

### CellSexID provides novel biological insights into macrophage ontogeny

The steady-state macrophage pool in adult skeletal has two principal origins: 1) a prenatal population established during muscle development from embryonic progenitor cells; and 2) a postnatal population derived from definitive monocytes recruited to the muscles [5]. The former macrophage population undergoes self-renewal in the tissue and is bone marrow-independent, whereas the latter population turns over rapidly and requires continuous replenishment by monocytes from the bone marrow. Accordingly, we next sought to determine whether CellSexID could ascertain the relative contributions of these two populations to macrophage subsets and their associated transcriptomic profiles in the main respiratory muscle, the diaphragm – where the role of macrophage ontogeny remains underexplored. To address this question, we generated additional chimeric mice in which CD45.1 male bone marrow was transplanted into CD45.2 female recipients, generating another “chimeric mouse diaphragm experimental dataset” (Figure 6a). Using our trained Random Forest model to predict cell origins from scRNA-seq analysis of sorted diaphragm macrophages at 8 weeks post-transplantation, the predicted percentages of donor (bone marrow-dependent) and recipient (bone marrow-independent) macrophages were 74% and 26%, respectively. This closely matched the percentages obtained from flow cytometry staining of CD45.1+ and CD45.2+ cells (70% and 30%, respectively) (Figure 6b). This confirms that CellSexID can be used to determine cellular origin in chimeric mice without the need for conventional tracking methods such as the CD45.1 allele or fluorescent marker genes, which are not readily available in most genetically modified mouse model strains. In addition to these label prediction, Supplementary Figure 11 visualizes the probability assigned to each cell by our classifiers. These visualizations provide an intuitive understanding of prediction uncertainty, showing how confidence varies across different regions of the embedding.

**Figure 6:**
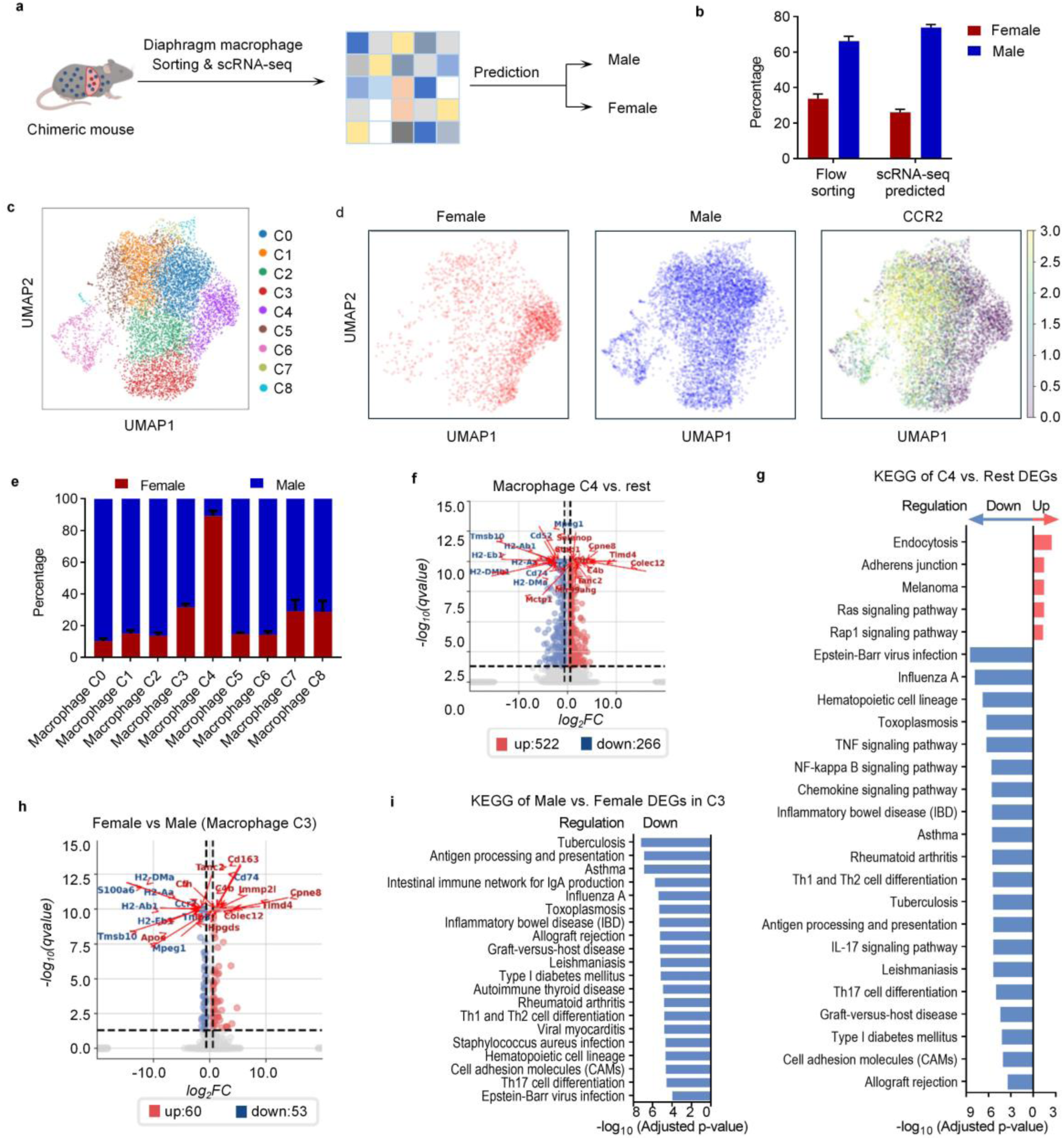
CellSexID enables origin-specific analysis of macrophage subpopulations, revealing unique gene expression profiles and functional pathways. **a**, Schematic overview of the experimental design. **b,** Proportional distribution of macrophages from recipient and donor origins in each mouse, determined by flow cytometry sorting and scRNA-seq predictions using our default classifier. Data represent eight biological replicates (n=8; mean ± SEM). **c**, Unsupervised clustering analysis identifies eight distinct macrophage groups, segregating cells from recipient (female) and donor (male) origins based on unique gene expression profiles. **d**, UMAP visualization showing the distribution of recipient- and donor-derived macrophages. Cell origins were annotated using our default classifier model and validated against the monocyte-derived macrophage marker *Ccr2*, supporting the accuracy of the predictions. **e,** Proportion of recipient- and donor-derived macrophages (annotated using our default classifier) within each identified cluster, highlighting origin-specific predominance in certain clusters (n=8; mean ± SEM). **f**, Volcano plot displaying the top 10 upregulated and downregulated genes in macrophage cluster C4 compared to all other clusters, highlighting significant gene expression differences. **g**, Pathway enrichment analysis of cluster C4 (top 20 pathways), indicating enhanced cell communication and proliferation alongside reduced immune and inflammatory responses compared to other clusters. **h**, Volcano plot highlighting genes with significant upregulation or downregulation in recipient-versus donor-derived macrophages within macrophage cluster C3. **i**, Pathway analysis of macrophage cluster C3 (top 20 pathways), revealing significantly downregulated immune-related pathways in recipient-derived macrophages compared to donor-derived macrophages.

We applied standard scRNA-seq data analysis pipeline for our preprocessed data according to common practice, which includes dimensionality reduction, clustering, and downstream analysis such as DEG and enrichment [64]. Visualizations such as UMAP and t-SNE can be incorporated once dimensionality reduction is finished. Our clustering leads to 9 distinct macrophage clusters in the diaphragm (visualized using UMAPs in Figure 6c). Please refer to Supplementary Figure 12 for our rationale in selecting the clustering resolution. Supplementary Figures 13 and 14 demonstrate the robustness of our clustering results, showing that a different resolution yields similar downstream analyses. Please refer to the Methods section for technical details. The sex origin of each macrophage cluster in the muscle was determined using the our default classifier, and there was a very high degree of overlap between male origin and expression of *Ccr2* (Figure 6d, Supplementary Figure 15), which is a marker for bone marrow-derived macrophages [65, 66]. All of the clusters were predominantly derived from donor (male) bone marrow monocytes with the exception of cluster C4, in which 89% of the macrophages were of recipient (female) origin (Figure 6e). To further assess the transcriptional consistency within this female-enriched cluster, we calculated a label score of 0.8041 based on local neighborhood sex label agreement, suggesting a high degree of local homogeneity in sex identity within this cluster. The donor bone marrow-dependent (male) macrophages displayed a dispersed and heterogeneous clustering pattern, consistent with a diverse population having varying gene expression profiles and functional states. In contrast, the bone marrow-independent (female) macrophages formed a tight, homogeneous cluster, indicating greater uniformity in their gene expression and most likely their function.

To gain further insights into the differential functions of bone marrow-dependent versus bone marrow-independent macrophages, we performed differential gene expression (DEG) analysis. In macrophage cluster C4, which is primarily composed of bone marrow-independent macrophages, 522 genes were significantly upregulated, and 266 genes were downregulated compared to all other clusters (Figure 6f). The top 10 upregulated genes included *Cfh, Colec12, Timd4, Mir99ahg, Selenop, C4b, Cpne8, Tanc2*, *Stab1*, and *Mctp1*, while the top 10 downregulated genes were *Cd74, H2-Eb1, H2-Ab1, H2-Aa, Ccr2, H2-DMa, Cd52, H2-DMb1, Mpeg1*, and *Tmsb10*. Pathway enrichment analysis revealed that the upregulated genes in cluster C4 were significantly associated with pathways such as “Endocytosis,” “Adherens Junction,” “Ras Signaling Pathway,” and “Rap1 Signaling Pathway” (Figure 6g). These pathways suggest enhanced cellular uptake processes, cell adhesion, and growth factor signaling, likely contributing to increased cell communication and proliferation. In contrast, the downregulated pathways included “Hematopoietic Cell Lineage,” “TNF Signaling Pathway,” and “NF-kappa B Signaling Pathway,” indicating reduced pro-inflammatory immune responses in this bone marrow-independent macrophage population. All of the macrophage clusters were predominantly composed of donor-derived (male) cells, except for cluster C4, in which 89% of the macrophages were of recipient (female) origin (Figure 6e). To further assess whether these transcriptional differences reflected biological divergence by origin rather than intrinsic sex effects, we compared gene expression between male- and female-classified cells across the dataset. We found a significant overlap (899 genes, *P* = 5.35 × 10⁻⁷⁹⁵) between the DEGs from the male vs. female comparison and those from the C4 vs. rest comparison (Supplementary Figure 16a), suggesting that CellSexID’s sex-based labeling faithfully captured origin-related identity. Importantly, since both donor and recipient macrophages are located within the same tissue microenvironment, they experience identical local cues, minimizing the influence of systemic sex differences. Therefore, the transcriptional divergence observed is more likely driven by ontogeny (donor vs. recipient origin) rather than by sex-specific regulation per se. Consistent with this interpretation, KEGG pathway analysis revealed that genes downregulated in recipient-derived (female) cells were enriched in immune-related pathways—such as antigen presentation, Th1/Th2 differentiation, and IBD—paralleling those identified in the cluster-level analysis (Figure 6g, Supplementary Figure 16b). Together, these findings support the conclusion that CellSexID’s sex-based predictions reflect meaningful biological variation rooted in cell origin.

To determine whether similar differences could be observed within a macrophage cluster containing cells from both origins, we applied our method, CellSexID, to macrophage cluster C3, which contains a mixture of donor-origin (68%) and recipient-origin (32%) macrophages. By enabling the identification of cellular origins within the same cluster, CellSexID allowed us to perform differential expression analysis between donor-derived (bone marrow-dependent) and recipient-derived (bone marrow-independent) macrophages. Without CellSexID, such comparisons would not have been possible, as the mixed origins of cells within the cluster obscure their distinctions. Using CellSexID, we identified 60 upregulated and 53 downregulated genes in recipient-origin macrophages compared to donor-origin macrophages (Figure 6h). Notably, pathway enrichment analysis revealed that the downregulated pathways in recipient-origin macrophages mirrored those observed in cluster C4, including pathways associated with antigen presentation, T-cell differentiation, and other pro-inflammatory immune responses (Figure 6i). These findings reinforce the distinction between bone marrow-dependent and bone marrow-independent macrophages and highlight the utility of CellSexID in enabling origin-specific analyses within mixed populations.

In addition, we examined transcriptional differences between cells of different origins—distinguished by their predicted sex labels (male donor vs. female recipient)—within the C3 cluster. DEG overlap analysis between these predicted-origin groups within C3 and the global male vs. female comparison revealed 192 shared genes (p = 7.53 × 10⁻²⁴², hypergeometric test) (Supplementary Figure 17), indicating a highly significant gene-level overlap. These results demonstrate that predicted sex can serve as a surrogate for studying cell origin–associated transcriptional programs, enabling insights that would be difficult to obtain in mixed-origin clusters like C3 without CellSexID.

## Discussion

This study addressed the critical challenge of tracking cell origins various biological context (e.g. chimeric models) by developing CellSexID, a machine learning-based tool that uses sex-specific gene expression in scRNA-seq data as a natural surrogate marker. By applying CellSexID to sex-mismatched chimeric models we have created a simple, low cost, and high-precision method that accurately distinguishes donor and recipient cells at single-cell resolution. Through careful feature selection, we identified 14 essential genes that include but are not limited to those located on sex chromosomes, which allow for robust sex prediction and thereby enable reliable cell origin tracking. CellSexID simplifies the cell tracking process by eliminating the need for biased labelling techniques dependent on exogenous marker genes, which commonly have variable levels of gene expression under different experimental conditions and can also lack cellular specificity (so-called “leaky expression”) [67, 68]. In addition, expensive and labor-intensive cross-breeding strategies are often required to incorporate exogenous cell origin markers into existing animal models [7, 8]. By taking advantage of the endogenous gene expression profile associated with biological sex, CellSexID is well positioned as an efficient, scalable solution in experimental as well as clinical settings.

There are four major contributions of CellSexID. **First**, the primary innovation of this study lies in its experimental design, which transforms the challenging task of cell origin tracking into a more manageable problem—sex prediction based on transcriptomic data. By utilizing sex-mismatched samples, machine learning models can efficiently identify informative gene features and classify cell origins using sex as a surrogate target, significantly reducing the costs and labor associated with traditional labeling methods. **Second**, our consensus-based committee feature selection pipeline robustly identifies minimal, yet highly informative gene marker sets across diverse datasets. Notably, this pipeline consistently selects reliable markers, demonstrating high predictive performance across datasets derived from various tissues and species. The default marker list includes both widely recognized sex-linked genes, such as *Xist*, *Ddx3y*, *Eif2s3y*, *Uty*, and *Kdm5d*, and, surprisingly, numerous highly predictive autosomal genes. The identification of autosomal markers, including *Uba52* (encodes a ubiquitin-ribosomal fusion protein that has been reported to play an important role in sex-related behavioral differences mediated by the amygdala [40]), *Lars2* (leucine-tRNA-synthase 2, was recently found to be highly enriched in endothelial cells of male mice [41]),and various ribosomal proteins (*Rpl* and *Rps*), challenges the conventional assumption that sex chromosome-linked genes predominantly determine sex prediction. Including autosomal markers led to improved predictive metrics, such as AUROC and AUPRC. These findings underscore the strength of CellSexID’s integrative machine learning framework in uncovering robust sex markers beyond traditional assumptions. Additionally, CellSexID offers flexibility by supporting feature selection on user-specified datasets and enabling users to input custom gene lists directly. **Third**, the general applicability of CellSexID extends beyond traditional animal chimeric models, as evidenced by successful validation in human datasets and clinically relevant contexts, including organ transplantation and sample demultiplexing. This broad applicability highlights the method’s potential utility across diverse experimental and clinical scenarios, where sex prediction serves effectively as a surrogate marker for cell origin tracking. **Finally**, using sex as a surrogate target for cell origin tracking enabled the discovery of novel biological insights into ontogeny-specific gene signatures and functional characteristics. Analyses performed on our chimeric mouse diaphragm experimental dataset illustrate these insights clearly. Thus, CellSexID provides a versatile and robust framework for generating biological discoveries across various tissues, species, and research contexts.

To demonstrate CellSexID’s real-world applicability, we validated its performance in chimeric mice with sex-mismatched bone marrow transplantation. In tracking the sex origin of macrophages in skeletal muscle using our tool, we were able to show high concordance with ground truth flow cytometry results, confirming its accuracy in distinguishing bone marrow-derived from bone marrow-independent cells. Furthermore, here we show the presence of important transcriptional differences in skeletal muscle macrophages based on cell origin (ie., between bone marrow HSC-derived and embryonic-derived macrophages), thus revealing how ontogeny helps to shape immune responses. Notably, we identified a unique subset of macrophages in the diaphragm which are overwhelming bone marrow-independent and display an anti-inflammatory and pro-regenerative gene expression profile. This discovery highlights CellSexID’s capacity to reveal ontogeny-driven functional differences within tissues where traditional methods have struggled to provide sufficient resolution. The characteristics of embryonic-derived tissue-resident macrophages are particularly relevant to various pathological states in which these cells are often lost from the diseased tissue [6, 69], and CellSexID provides a powerful new tool for tracking their behavior at different stages of pathology.

A limitation of CellSexID is its reliance on sex differences between donor and recipient animals, which could potentially influence cell states or functions in certain contexts. While this dependency does not affect the main goals of this study, it may introduce bias in specific cell types where sex differences play an important role in determining phenotype. To address this limitation, future studies could implement bi-directional chimeric experiments with both male and female donors and recipients. This approach not only strengthens the robustness of our method but could also promote a greater understanding of the more nuanced aspects of sex differences in biological systems.

CellSexID has the potential to transform research by providing a precise, scalable, and cost-effective tool for tracking cell origins at single-cell resolution. By leveraging endogenous sex-specific gene expression, it eliminates the need for exogenous labeling or genetic engineering, overcoming limitations such as high costs and technical complexity in traditional methods. This innovation is broadly applicable across diverse research settings and addresses a critical gap in experimental biology. By enabling high-resolution tracking of cellular dynamics, CellSexID empowers researchers to explore ontogeny-driven differences in cell function across contexts such as organ transplantation, single-cell demultiplexing, single-cell dataset quality control and contamination identification, retrospective analysis of public datasets lacking sex annotation, sex-associated disease mechanisms investigation, immune regulation, tissue repair, and developmental biology. Our analysis on the chimeric mouse diaphragm experimental dataset demonstrates CellSexID’s ability to identify functionally distinct cell populations, such as embryonic-derived macrophages with pro-regenerative properties, underscores its utility in uncovering fundamental biological insights. By providing a robust framework for understanding cellular dynamics, it holds great promise for advancing biomedical research and enabling the development of targeted therapeutic strategies.

## Methods

### Experimental animals

Experiments utilized mice carrying CD45.2 and CD45.1 alleles, obtained from The Jackson Laboratories in Bar Harbor, ME, USA. The animals were housed under controlled conditions in the vivarium of the McGill University Health Centre, following the standards of the Canadian Council on Animal Care, the McGill University Animal Ethics Committee, and ARRIVE guidelines. Before starting the study, the mice underwent a 7-day acclimation period in our facility. Skilled personnel provided high-quality care, including regular checks, to ensure the animals’ well-being throughout the research.

### Bone marrow transplantation

Chimeric mice were created following our previously outlined protocol [59]. In brief, 6-week-old female CD45.2 mice received whole-body X-ray irradiation using the X-RAD SmART irradiator (Precision X-ray, USA). Lead shielding was applied to the diaphragm area to protect resident macrophages. The irradiation was fractionated into two doses of 6 Gy, delivered 4 hours apart, with settings of 225kV, 13mA, and 1.0265 Gy/min. Six hours after the final irradiation, an intraperitoneal injection of 30 mg/kg busulfan (Sigma, USA) was administered to eliminate any remaining bone marrow protected by diaphragm shielding. Twenty-four hours post-irradiation, CD45.2 mice received an intravenous injection of 2x10^7^ bone marrow cells from age-matched CD45.1 male donors. To reduce infection risk, the recipient mice were provided with enrofloxacin in their drinking water for 7 days following irradiation. An 8-week recovery period was then allowed to ensure complete bone marrow reconstitution. Two female mice were used as recipients for the chimeric mouse diaphragm validation dataset and eight female mice were used as recipients for the chimeric mouse diaphragm experimental dataset.

### Flow cytometry

To quantify and identify macrophages in diaphragm, flow cytometry was performed as described in prior studies [59, 70]. Mice were anesthetized using isoflurane before euthanasia, followed by heart perfusion with 20 ml of phosphate-buffered saline (PBS) (Wisent, Cat. #811-425), with an additional 20 ml administered after cutting the dorsal aorta. The diaphragm tissue was dissected, minced, and digested in PBS containing 0.2% collagenase B (Roche, Cat. #11088815001) for 1.5 hours to create a cell suspension. Diaphragm cells were stained with a viability dye to distinguish live from dead cells. For surface marker staining, cells were incubated at 4°C in the dark with primary antibodies, including anti-CD45.1 (clone A20, Cat. #553776), anti-CD45.2 (clone 104, Cat. #562129), anti-SiglecF (clone E50-2440, Cat. #562757), and anti-CD11c (clone HL3, Cat. #564986) (all from BD Biosciences, NJ, USA); and anti-CD11b (clone M1/70, Cat. #101226) and anti-F4/80 (clone BM8, Cat. #123114) (both from Biolegend, CA, USA). To prevent non-specific binding, cells were treated with Fc blocking solution (anti-CD16/CD32, clone Ab93, BD Biosciences, Cat. #567021). To identify diaphragm macrophages, we defined them as cells positive for CD45 (either CD45.1 or CD45.2), CD11b and F4/80, and negative for SiglecF and CD11c. For the model validation experiment, we specifically added Cell Multiplexing Oligo (CMO, 10X Genomics, cat. # 1000261) to sorted populations of CD45.1 and CD45.2 macrophages to differentiate between donor and recipient mouse macrophages. In experiments addressing our research question using the model, CMOs were added to the unstained muscle cell suspension before sorting to distinguish cells originating from different mice. All macrophages were sorted in a PBS solution containing 20% fetal bovine serum (FBS) to maintain cell viability and integrity throughout the process.

### Chimeric mouse diaphragm validation dataset

For the validation experiment, we utilized a ratio of 80% Cell Multiplexing Oligo (CMO)-labelled CD45.1 macrophages to 20% CMO-labelled CD45.2 macrophages. This mixture was loaded onto a 10x Genomics Chip G (Cat. #1000127) and processed using the 10x Chromium Controller, following the manufacturer’s instructions.

### Chimeric mouse diaphragm experimental dataset

For the experiments aimed at addressing the research question, a mix of CD45.1 and CD45.2 positive cells was added to the chip without distinguishing between the two types. The library preparation was then conducted in accordance with 10x Genomics protocol CG000388.

All libraries were sequenced by Illumina NovaSeq6000 S4 v1.5.

### Single-cell data acquisition

For our generated single-cell data (chimeric mouse diaphragm validation and chimeric mouse diaphragm experimental datasets), at 6 weeks post-transplantation, we used the same bone marrow transplantation protocol as described for generating chimeric mice. Wild-type (WT) female recipient mice were transplanted with bone marrow cells from WT male donors. Following the transplantation protocol, the mice were allowed to recover for 8 weeks to ensure complete bone marrow reconstitution. Subsequently, macrophages from the diaphragms of these mice were isolated via cell sorting, specifically targeting populations expressing CD45+ and F4/80+ markers to ensure high purity. Single-cell RNA sequencing was then performed on these isolated macrophages. Sequencing reads were processed using 10x Genomics Cell Ranger 7.1.0. Each library was processed using the Cell Ranger [71] multi pipeline. Specifically, we used the reference genome refdata-gex-mm10-2020-A with a minimum assignment confidence of 0.7. The libraries included gene expression and multiplexing capture data. The samples analyzed were WT_to_WT samples. Using Cell Ranger multi, each library was aligned to an indexed mm10 genome, generating an aggregated matrix of WT cells. This aggregation normalized the number of confidently mapped reads per cell in each library, preparing the data for downstream analysis.

### Single-cell data preprocessing

The SCANPY [72] (version 1.9.5) was applied to preprocess single-cell raw count matrices for all datasets. For GSM6153751 (female mice cells) and GSM6153750 (male mice cells) from adrenal gland dataset, we applied minimum cell expression threshold of 3, followed by log transformation and normalization. Since this served as the major gene selection dataset, we retained all genes without gene filtering to preserve comprehensive gene expression information. For the chimeric mouse diaphragm validation dataset, we used the same processing pipeline but applied a gene threshold of 100 and doublet removal using Scrublet to exclude potential multiplets. As this was a relatively small dataset, this approach removed only 3 cells, resulting in 1649 cells. For all cross-tissue validation datasets including mouse kidney (GSE129798), mouse hypothalamus-PVN (GSE201032), mouse colon (GSM2967048), and human ATL blood (GSE294224), human AML BM(GSE289435), and human thymus (GSE262749) datasets, we applied stringent quality control parameters: minimum cell expression threshold of 3, removing cells expressing fewer than 200 genes, and mitochondrial gene percentage threshold of 5%, followed by log transformation and normalization to a total count of 10,000 per cell. For the kidney transplant scRNA-seq data (GSE151671), we applied the same parameters except for the mitochondrial gene percentage threshold, as applying the 5% threshold would result in loss of nearly all cells due to the inherently high mitochondrial gene expression in transplant tissue samples, likely reflecting cellular stress and metabolic alterations associated with transplantation procedures. For the chimeric mouse diaphragm experimental dataset, we first performed quality control filtering genes not expressed in at least three cells. We filtered cells expressing more than 6200 genes or fewer than 200 genes. Cells with more than 20% mitochondrial reads were regarded as dead cells and also filtered. Immunocytes were specifically filtered by selecting cells expressing *Ptprc*. We also performed doublet removal using Scrublet [73] to exclude potential multiplets. Subsequently, the single-cell data matrix was normalized to a total count of 10,000 per cell and log1p-transformed. Then, for cell origin classification, only the sex markers’ expression was used for model input. For clustering analysis, highly variable genes were detected using SCANPY’s tool ’pp.highly_variable_genes’ with default parameters, resulting in 1895 genes. The effects of total counts and mitochondrial gene percentage were regressed out. The data was then scaled, capping the values at 10.

### Classifier committee

We employed a classifier committee to predict sex based on gene expression data. The committee consists of four distinct machine learning classifiers, each contributing unique perspectives to the task. By working together, the committee helps to identify the most relevant features for maximizing prediction accuracy. While the best-performing classifier is ultimately chosen to make the final prediction on the Chimeric mouse diaphragm test dataset, the combined efforts of the committee enhance the overall robustness and effectiveness of the feature selection process.

The choices of the models are based on their capabilities in feature selection and dealing with our classification tasks. **XGBoost Classifier** is a highly scalable model grounded in gradient boosting frameworks and is capable of capturing complex nonlinear patterns [74, 75]. It evaluates feature importance through a combination of metrics that describe how feature enhances accuracy and decision-making, which includes Gain – the improvement in accuracy a feature provides to the decision points it influences, Cover – the proportion of data points a feature impacts, and Frequency – how often a feature is utilized across the ensemble of decision trees. **Random Forest Classifier [76]** leverages an ensemble of decision trees and quantifies feature importance through the mean decrease in impurity, often expressed in terms of Gini impurity or entropy. This approach calculates the extent to which each gene reduces uncertainty across trees, with a higher reduction indicating a stronger influence on model predictions. This method is particularly effective in discerning the genes that consistently refine the model’s outputs by creating clearer, more defined paths in the decision process. For the **Support Vector Machines (SVM)** [77–79] with a linear kernel used in our method, feature importance is derived directly from the model’s coefficients. These coefficients are integral to the hyperplane that separates the classes in the dataset. Each coefficient quantifies the influence of a corresponding gene on the classification boundary, with its magnitude indicating the strength of the impact and its sign denoting the direction (positive or negative). This linear approach provides a clear and quantifiable measure of each gene’s contribution to the model’s decision-making process, facilitating the identification of key genetic features crucial for predictive accuracy. Linear SVMs are preferred for their ability to perform well in high-dimensional spaces with clear margins of separation, and their simplicity and effectiveness in binary classification scenarios. Lastly, **Logistic Regression** [80, 81]determines the importance of genes through its coefficients within a linear context, where the magnitude of each coefficient indicates the extent to which a gene affects the model’s output. Larger absolute values suggest a stronger influence, making this approach straightforward for gauging the impact of individual genes on the prediction of the target variable. Logistic Regression is favored for its interpretability and the provision of probability scores for outcomes, which can be crucial for understanding risk factors in medical or biological studies. These methods, grounded in robust statistical and mathematical principles, allow for a detailed exploration of gene significance in predictive modeling. In our implementation, Logistic Regression, Random Forest and Support Vector Machine are implemented in Python packages Scikit-learn [82] and XGBoost is implemented using package ‘XGBoost’ [14].

### Sex-relevant feature selection

We used the public mouse adrenal gland dataset to select the most significant gene features for predicting the sex of individual cells. First, each model in the committee is trained on the public adrenal gland dataset using all gene features and employed the built-in feature selection functions provided by their respective libraries to compute feature importance scores. The top 𝐾(default 20, see justification of this hyperparameter in Supplementary Figure 2) important features are selected. Then features selected by at least 𝐶 committee members are used as final selected features. We set 𝐶 to be 3 out of 4 (75%) to claim a majority and Supplementary Figure 4 justifies our choice. For any gene feature 𝑔, it will be selected in the final feature set 𝑮_𝐟𝐢𝐧𝐚𝐥_ if and only if it is selected by at least three models.

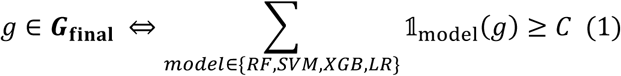

Where each 𝟙_model_(𝑔) represents an indicator function indicating if the gene feature is selected by a specific model (SVM: Support Vector Machine; LR: Logistic Regression; RF: Random Forest; XGB: XGBoost).

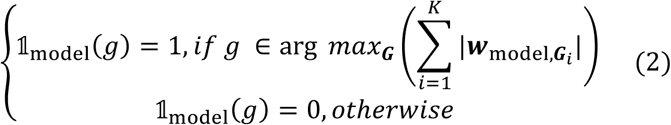

Different 𝒘 corresponds to different models’ feature importance score returned by the Python library after training using all gene features on the public mouse adrenal gland dataset. |𝒘_𝑆𝑉𝑀_| represents the absolute value of the trained SVM’s weights. |𝒘_𝐿𝑅_| denotes the absolute value of the trained Logistic Regression classifier’s coefficients. 𝒘_𝑅𝐹_ is the importance of the trained Random Forest derived from

the mean decrease in impurity across all trees in the forest. 𝒘_𝑋𝐺𝐵_ is the importance of the score of trained XGBoost derived from the Gain, Cover, and Frequency as mentioned in the previous paragraph. The subscription 𝑮_𝑖_ specifies the gene that the weight corresponds to.

The model committee also computes a scaled feature importance score for each gene feature, designed to reflect consensus among the committee members. This score is determined as follows: first, each committee member’s score is normalized to a range between 0 and 1. The normalized scores are then summed across all committee members. To further refine the score, a bonus of 0.25 is added for each committee member that selects the feature, while a penalty of 0.25 is subtracted for each committee member that does not select the feature. As a result, the final bonus/penalty component of the score will range between -1 (if none of the four committee members select the feature) and +1 (if all four committee members select the feature), providing a balanced measure of agreement across the committee. All the gene features will be ranked based on their feature selection scores. Once the feature selection process is completed, we use Random Forest as the classifier to predict sex by default.

### Sex classifier evaluation

Using the selected 14 final gene features, we tested the same models (Logistic Regression, Random Forest, Support Vector Machine, XGBoost) both on the mouse adrenal datasets and the chimeric mouse diaphragm validation dataset. Based on the selected sex-linked markers, classifiers output the probability of belonging to each class, which is converted to class binary prediction by setting a threshold, a common practice implemented in Scikit-learn. The predicted labels are subsequently used for evaluations. For testing on the mouse adrenal dataset, we used 5-fold cross validation to compute Accuracy and F1 score. Standard deviations are shown as error bars. For calculating Area Under the Receiver Operating Characteristic curve (AUROC) and the Area Under the Precision-Recall curve (AUPRC) we used functions in scikit-learn. For testing the model on the chimeric mouse diaphragm macrophage dataset, the full public dataset was used for training the model. For generating the bar plots in Figure 4 e and f, AUPRC and AUROC are calculated using bootstrapping with 100 iterations. In each iteration, the adrenal gland training dataset is randomly resampled with replacement to get the same number of samples as the original training set, but the test set (chimeric mouse diaphragm validation dataset) is unchanged, ensuring a consistent basis for performance comparison.

These metrics, along with their 95% confidence intervals derived from the bootstrap samples, provided comprehensive insights into the models’ predictive capabilities. While all models demonstrated excellent performance, we selected Random Forest as our default classifier for CellSexID. However, all four classifiers achieved comparable results and can be used interchangeably. The Random Forest classifier was subsequently used for annotating the cells in the single-cell dataset from our experiments.

### UMAP and clustering analysis for the single-cell data

Principal Component Analysis (PCA) [83] was performed with 50 components. For the chimeric mouse diaphragm experimental dataset, Harmony [84, 85] [86] was applied to remove batch effects across 8 samples and adjust the PCA basis. The neighborhood graph of cells was computed using the PCA representation using the SCANPY’s tool ‘pp.neighbors’ by considering 10 nearest neighbors and using the first 40 principal components, followed by UMAP [87] embedding using the ‘tl.umap’ tool with default parameters and Leiden clustering using SCANPY’s ‘tl.leiden’ tool with a resolution of 0.6. After further filtering out clusters with fewer than 200 cells, 9 macrophage clusters were identified. The choice of clustering resolution is based on Silhouette scores. In Supplementary **UMAP and clustering analysis for the single-cell data:** Principal Component Analysis (PCA) [83, 84] was performed with 50 components. For the chimeric mouse diaphragm experimental dataset, Harmony [85, 86] was applied to remove batch effects across 8 samples and adjust the PCA basis. The neighborhood graph of cells was computed using the PCA representation using the SCANPY’s tool ‘pp.neighbors’ by considering 10 nearest neighbors and using the first 40 principal components, followed by UMAP [87] embedding using the ‘tl.umap’ tool with default parameters and Leiden clustering using SCANPY’s ‘tl.leiden’ tool with a resolution of 0.6. After further filtering out clusters with fewer than 200 cells, 9 macrophage clusters were identified. The choice of clustering resolution is based on Silhouette scores. In Supplementary Figure 12 we show Silhouette scores [88] for different resolutions. Based on the plot, we select the resolution 0.6, which is before significant score drop. In addition, to justify the robustness of our analysis, we performed the same set of analysis as in Figure 6 using resolution 0.4 in Supplementary Figure 13, which shows very similar results. In Supplementary Figure 13, in the C3 vs. rest comparison, shared genes such as *Timd4* and *Tmab10*, as well as pathways including NF-kappa B signaling and TNF signaling pathway, show strong similarity to the results at resolution 0.6. In the C1 female vs. male comparison, consistent genes like *Tmsb10*, *H2-DMb1*, and *Timd4,* together with pathways such as antigen processing and presentation and Th1 and Th2 cell differentiation, also indicate overlap with the resolution 0.6 analysis. Furthermore, we also compared the differentially expressed genes (DEGs) between corresponding clusters at different resolutions, shown in Supplementary Figure 14. These similarities justify our analysis results in the Figure 6 are robust to the choice of clustering resolution.

### Enrichment analysis

Differentially expressed genes (DEGs) were identified using SCANPY’s tool ‘tl.rank_genes_groups’ with the Wilcoxon rank-sum test method [89]. Sex-related genes were removed from the analysis. Batch correction was not applied because our analysis shows it removes biological signal and does not affect our analysis results much. To evaluate the impact of batch correction on downstream analysis, we applied Scanorama to correct the gene expression matrix of the chimeric mouse diaphragm validation dataset. However, this substantially reduced sex-based signals critical to our study. For instance, the AUROC of the Random Forest classifier dropped from 0.837 (uncorrected) to 0.607 (corrected) (Supplementary Figure 18 a–b), indicating that batch correction attenuated meaningful biological variation. This effect has been noted in prior studies (Xiong et al. [90], and others have reported that batch correction may inadvertently remove genuine biological variation—especially when the variable of interest (e.g., sex) is partially confounded with batch structure [91]. To preserve sex-specific signals, batch correction was not applied to the gene expression matrix in our final pipeline. In the DEG analysis, up-regulated genes were selected with Benjamini-Hochberg adjusted p-values less than 0.05 and log2 fold changes greater than 0.6, while down-regulated genes were selected with Benjamini-Hochberg adjusted p-values less than 0.05 and log2 fold changes smaller than -0.6. Top regulated genes were visualized through volcano plots, generated using the custom scatter plot function implemented in Python tool Matplotlib. If the number of top regulated genes exceeded 200, the top 200 genes were used; if fewer than 200 top regulated genes were identified, all top genes were used. The top up- and down-regulated genes were then subjected to KEGG 2019 Mouse [92–95] enrichment analyses using the GSEApy [96] tool ‘enrich’ with KEGG for enrichment analysis.

## Data availability

https://www.ncbi.nlm.nih.gov/geo/query/acc.cgi?acc=GSM6153751 https://www.ncbi.nlm.nih.gov/geo/query/acc.cgi?acc=GSM6153750

The public single-cell RNA-sequencing datasets used in this study are available from the Gene Expression Omnibus (GEO). The primary training dataset consists of adrenal gland CD45⁺ immune cells from 7-week-old C57BL/6 mice, originally published by Dolfi et al. [28], and can be accessed under accession numbers GSM6153751 for female mice (3,963 cells) and GSM6153750 for male mice (2,685 cells). For cross-tissue validation, additional mouse datasets were included: kidney data (GSE129798) [54], hypothalamic paraventricular nucleus (PVN) data (GSE201032) [55], and colon data from the Tabula Muris project (GSM2967048) [56]. The colon dataset served a dual purpose as both an independent test set and a training source for marker-based models. Human validation datasets included dendritic cell-like transitions in peripheral blood mononuclear cells (PBMCs) from T-cell leukemia–lymphoma (ATL) patients (GSE294224) [57], bone marrow mononuclear cells from acute myeloid leukemia (AML) patients (GSE289435) [33], thymic mimetic epithelial cells (GSE262749) [58], and kidney transplant immune cell data (GSE151671) [63]. Among these, the bone marrow dataset functioned as the primary training source for human transfer learning experiments. All datasets are accessible through the GEO portal at https://www.ncbi.nlm.nih.gov/geo. The chimeric mouse diaphragm macrophage single-cell datasets experimentally generated in this study will be made available upon acceptance.

## Code availability

Code for the models and results reproduction is publicly available on GitHub: https://github.com/mcgilldinglab/CellSexID

## Acknowledgement

This work was funded by grants awarded to JD and BJP. We gratefully acknowledge the support from the Canadian Institutes of Health Research (CIHR) under Grant Nos. PJT-180505 and 180644; the Funds de recherche du Québec - Santé (FRQS) under Grant Nos. 295298 and 295299; the Natural Sciences and Engineering Research Council of Canada (NSERC) under Grant No. RGPIN2022-04399; and the Meakins-Christie Chair in Respiratory Research.

