## Supplementary Figures for "Computational Tracking of Cell Origins Using CellSexID from Single-Cell Transcriptomes"

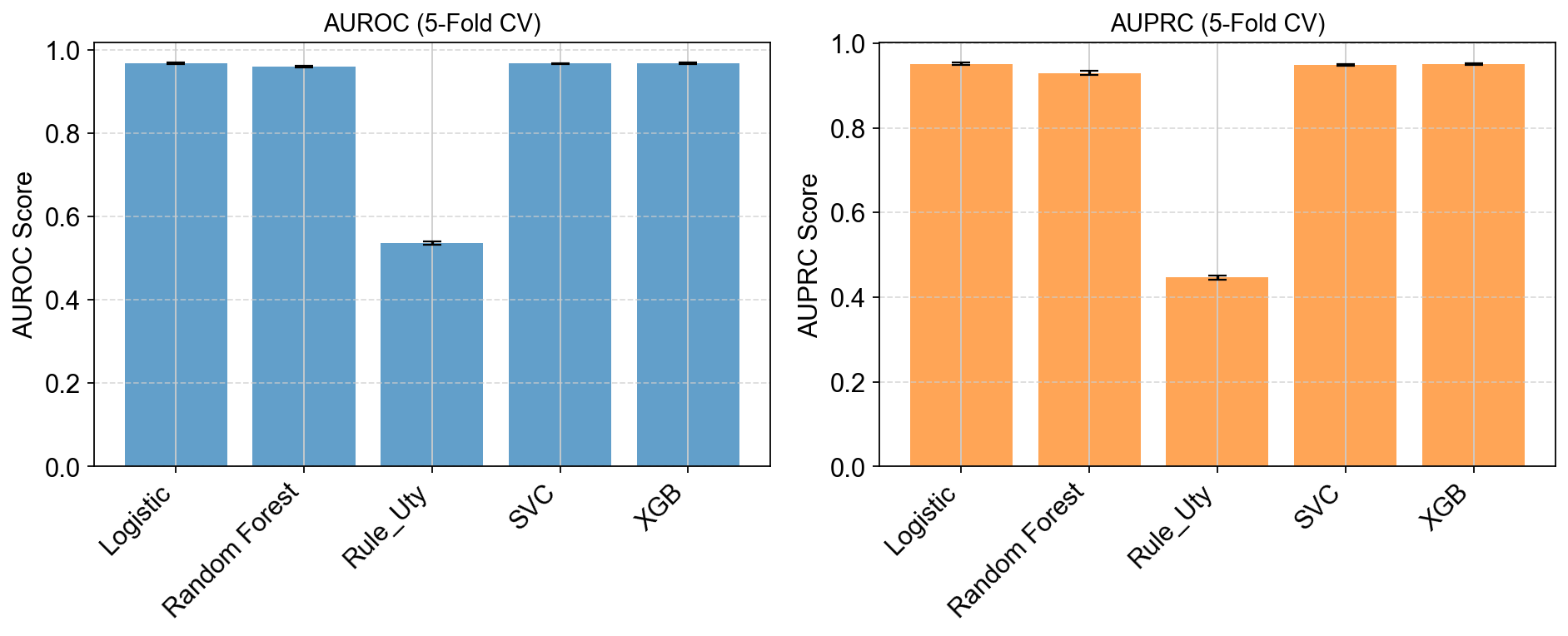

**Supplementary Figure 1: Bar plots comparing machine learning models’ performance evaluated using AUROC and AUPRC on the chimeric mouse diaphragm validation dataset.** 5-fold cross-validation is used for evaluation. The four models adopted by CellSexID are trained on the

full mouse adrenal gland

dataset. They show superior performance than the rule-based baseline method.

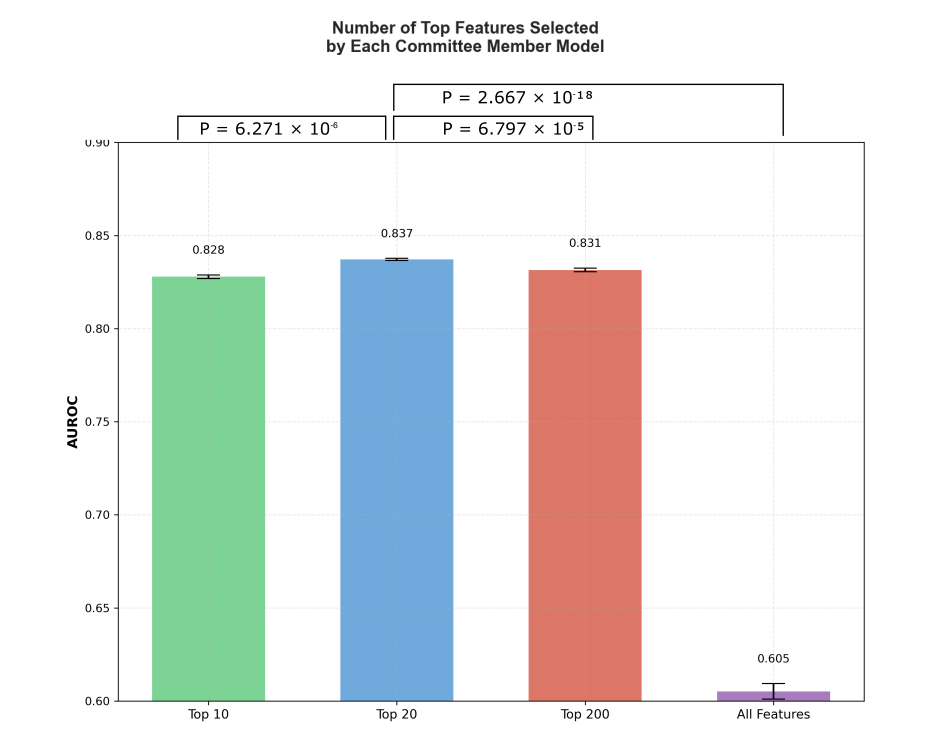

**Supplementary Figure 2**: **Justification for selecting the top 20 genes in committee-based feature selection.**

AUROC of Random Forest (RF) classifier performance across 15 random seeds, trained on the mouse adrenal gland dataset and tested on the chimeric mouse diaphragm validation dataset, using different numbers of top features (top 10, 20, 200 per committee member, and all features) selected during the committee-based feature selection process. Bars indicate the mean AUROC across 15 replicates; error bars show standard deviation. Using the top 20 genes as a feature selection hyperparameter resulted in significantly better performance than using the top 10 (one-tail t-test p-value = 6.271 × 10⁻⁶), top 200 (one-tail t-test p-value = 6.797 × 10⁻⁵), or all features (one-tail t-test p-value = 2.667 × 10⁻¹⁸). The top-20 setting was chosen to balance predictive performance and feature parsimony, retaining the most informative gene features while reducing the number of selected genes.

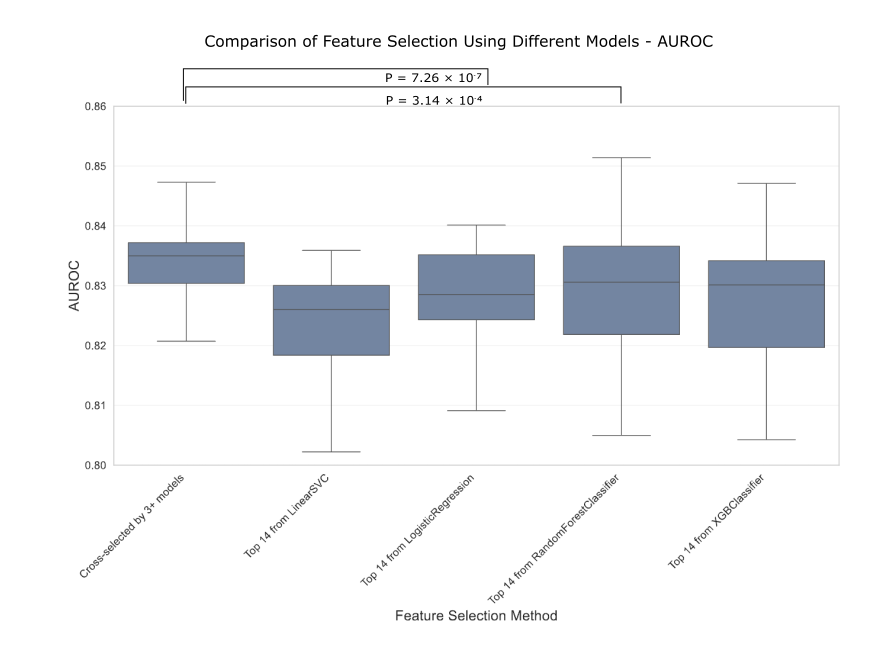

**Supplementary Figure 3: Justification of the committee feature selection strategy.** Box plots comparing the performance of different feature selection methods. The x-axis shows feature sets selected by individual models (Logistic Regression, Linear SVC, XGBoost, Random Forest) or by committee consensus (cross-selected by 3+ models). The y-axis shows AUROC values from Scikit-learn’s MLP classifiers tested on the chimeric mouse diaphragm validation dataset. All models were trained on the adrenal gland dataset using 14 features selected by each respective method. Results are shown for 15 random seeds where cross-selected features achieved the highest mean AUROC performance. Each box represents 75 data points (5 models × 15 seeds). The committee selection demonstrates comparable or superior performance to individual model selections using one tailed t-test. The committee approach provides a more robust and generalizable feature set that reduces the risk of overfitting to any single model's selection bias.

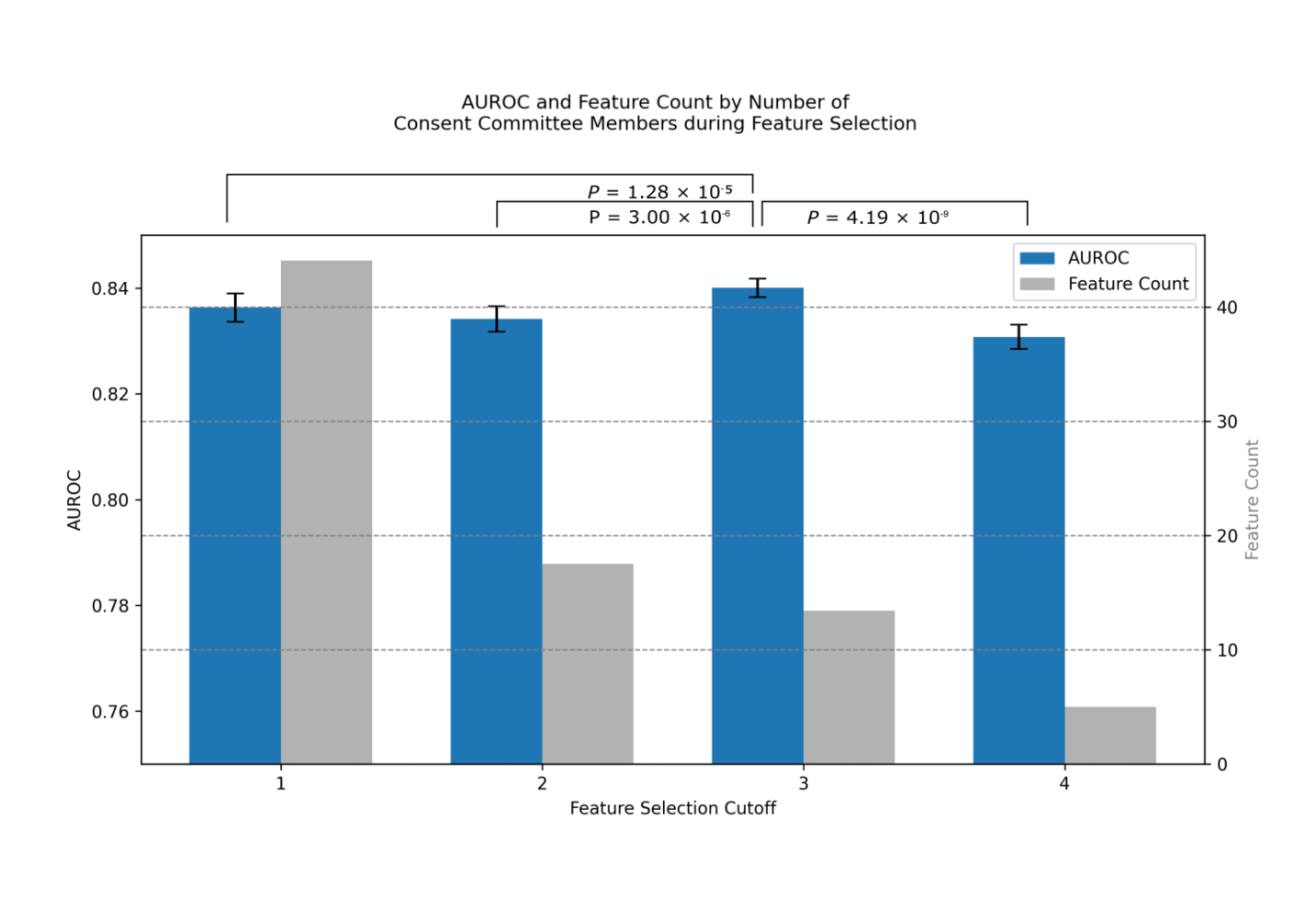

**Supplementary Figure 4**: **Justification of the number of consent committee members in the feature selection.** The x-axis represents feature selection stringency, showing models using features selected by at least C = 1, 2, 3, or 4 committee members. The mouse adrenal gland dataset was used for training, and the chimeric mouse diaphragm validation dataset for testing. The primary y-axis (left, blue bars) shows the Area Under the Receiver Operating Characteristic curve (AUROC) with error bars indicating standard deviation across five random seeds and 4 models. The secondary y-axis (right, grey bars) displays the corresponding number of selected features. Feature counts: 44 markers for cutoff 1, 18 markers for cutoff 2, 14 markers for cutoff 3, and 5 markers for cutoff 4. Statistical analysis using one-tailed t-tests revealed that cutoff 3 achieved significantly higher AUROC compared to all other cutoffs: vs. cutoff 1 (*p* = 1.28 × 10⁻⁵), vs. cutoff 2 (*p* = 3.00 × 10⁻⁶), and vs. cutoff 4 (*p* = 4.19 × 10⁻⁹). Based on these results, we selected cutoff 3 as optimal, resulting in the highest predictive performance (AUROC = 0.840) with a feature set of 14 markers.

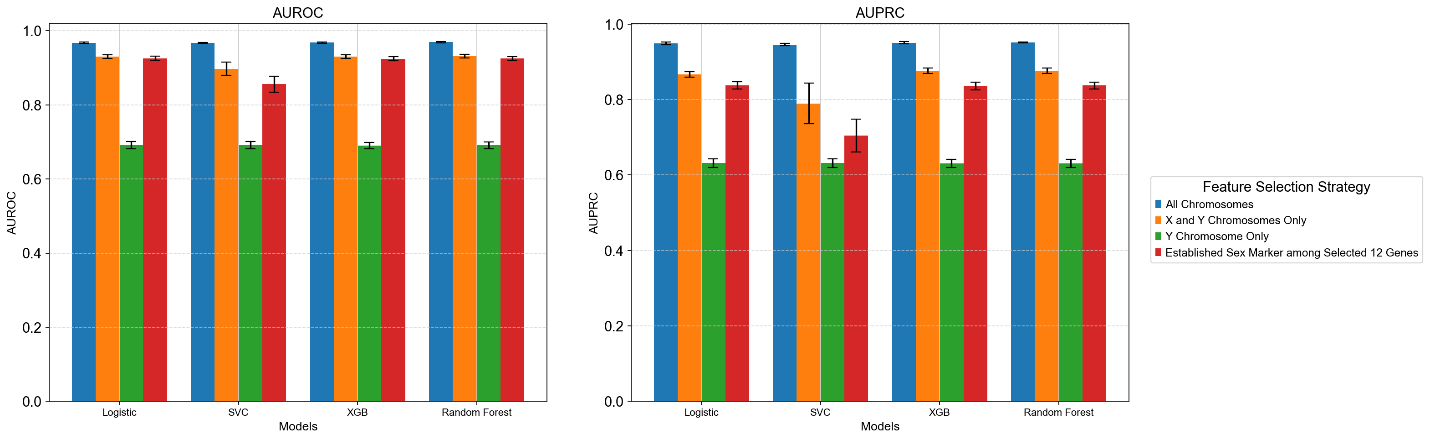

**Supplementary Figure 5**: **Comparison of model performance across different gene selection strategies.** Performance metrics (AUROC and AUPRC) were evaluated by results from 5-fold cross-validation for four different models (Logistic Regression, SVC, XGBoost, and Random Forest) using mouse adrenal gland dataset across three feature strategies: applying feature selection pipeline on genes from all chromosomes (blue), on selected X chromosome genes (including key genes involved in X-inactivation and dosage compensation like *Xist, Tsix, and Rnf12*) together with Y chromosome genes (orange), on only Y chromosome genes (green), and on established sex markers that are among our 14 genes (*Xist, Ddx3y, Eif2s3y, Uty, and Kdm5d*, shown in red). Error bars represent standard deviation across five folds. The results demonstrate that models trained using features from all chromosomes consistently outperform those using only sex-linked genes, with Y chromosome-only features showing the lowest performance across all models. This analysis justifies the usefulness of autosomal genes and our decision to include genes from all chromosomes in our final model rather than restricting it to sex-linked genes only.

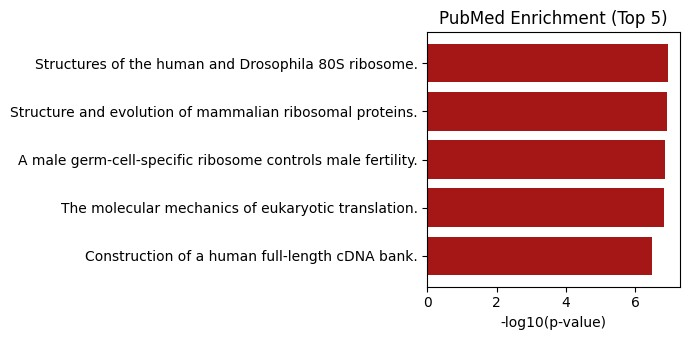

**Supplementary Figure 6: Functional enrichment analysis of the autosomal genes in the mouse sex-linked marker list.** Topgene’s PubMed gene sets are used for the analysis. Statistically significant terms show relevance to sex difference, such as male germ cell-specific ribosome and translation-related pathways, which are functionally associated with sex-biased expression and male fertility.

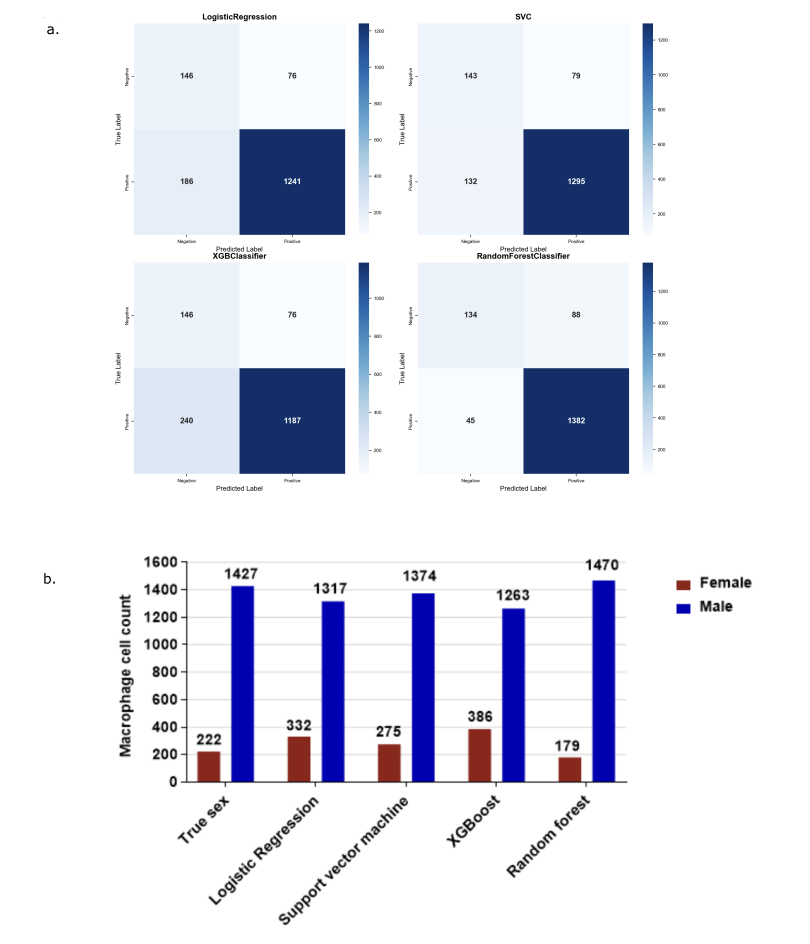

**Supplementary Figure 7: Performance of Random Forest, Logistic Regression, XGBoost, and SVM classifiers on the chimeric mouse diaphragm validation dataset.** **a**, Confusion matrices for all classifiers based on the selected gene features, demonstrating consistently strong predictive accuracy for cell origin comparable to our default classifier Random Forest (see Figure 4b). **b,** Comparison of the predicted sex (cell origin) ratios obtained from all classifiers with the ground truth, indicating all models accurately predict cell origin. These analyses provide comprehensive evidence supporting the robustness of all four classifiers in the CellSexID pipeline.

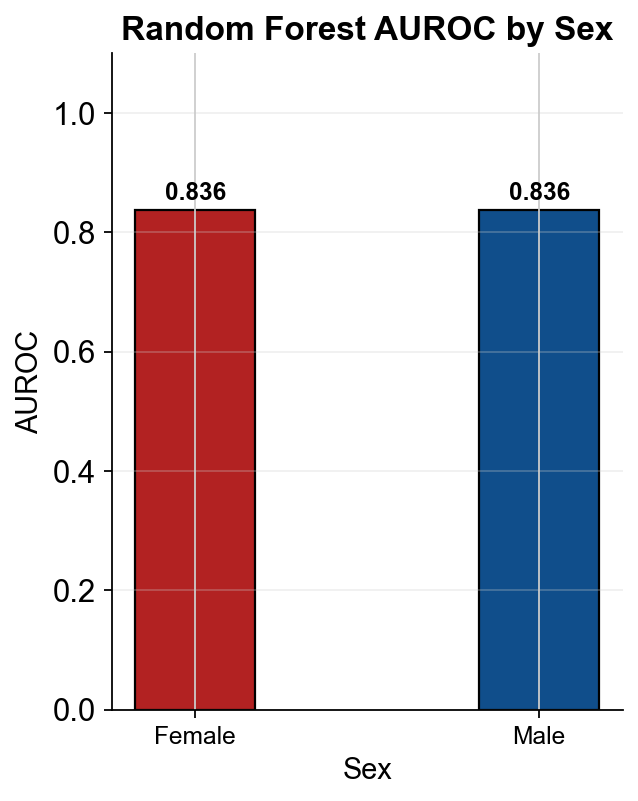

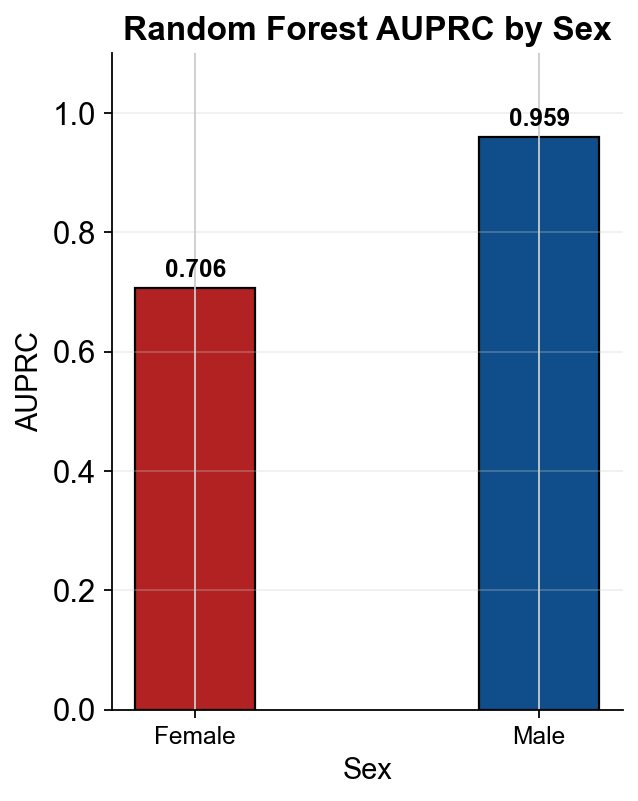

**Supplementary Figure 8: Class-specific prediction performance metrics.** Class-specific prediction performance metrics of our default classifier, trained on the adrenal gland dataset and tested on our chimeric mouse diaphragm validation dataset (linked to Figure 4b). AUROC and AUPRC of each sex’s prediction are shown.

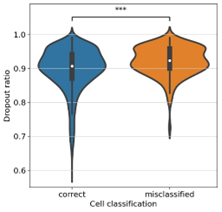

**Supplementary Figure 9:** **Comparison of dropout ratios between correctly classified and misclassified cells in the chimeric mouse diaphragm validation dataset.** Violin plots showing the distribution of dropout ratios (fraction of genes with zero expression) in correctly classified versus misclassified cells. Misclassified cells exhibit significantly higher dropout ratios (*p* = 8.5066 × 10⁻⁵, Mann–Whitney U test), indicating increased technical noise or sparsity, which may contribute to classification errors.

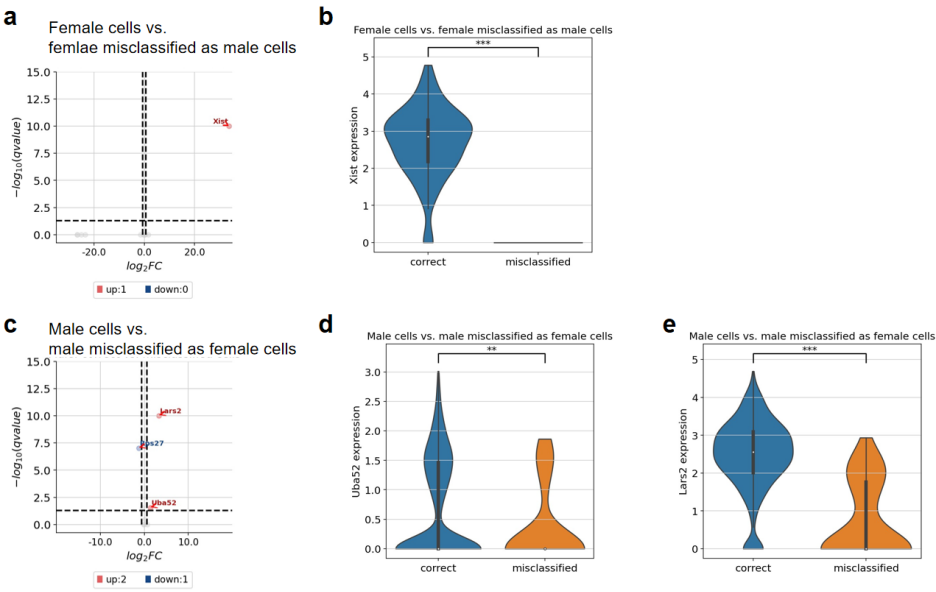

**Supplementary Figure 10:** **Differential expression analysis and violin plots showing expression of sex-related genes in correctly classified and misclassified cells in the chimeric mouse diaphragm validation dataset**. **a,** Volcano plot of correctly classified female cells vs. female cells misclassified as male cells. **b,** Violin plot of *Xist* expression in female cells, showing significantly higher expression in correctly classified cells compared to misclassified cells (Mann-Whitney U test, *p* = 6.23 × 10⁻³⁶). **c,** Volcano plot of correctly classified male cells vs. male cells misclassified as female cells. **d,** Violin plot of *Uba52* expression in male cells, showing higher expression in correctly classified cells (Mann-Whitney U test, *p* = 3.74 × 10⁻³). **e,** Violin plot of *Lars2* expression in male cells, showing significantly higher expression in correctly classified cells (Mann-Whitney U test, *p* = 4.74 × 10⁻¹⁸).

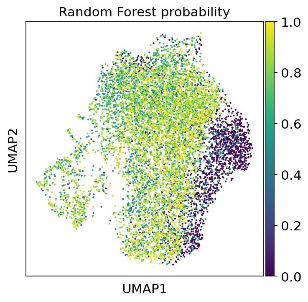

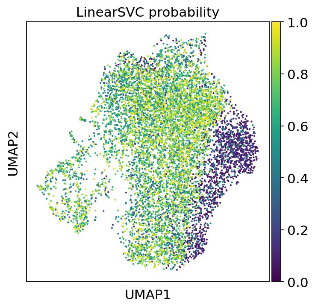

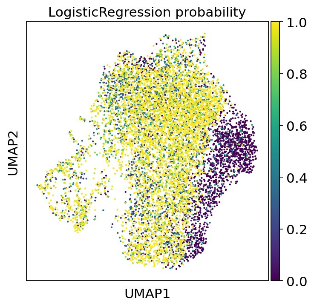

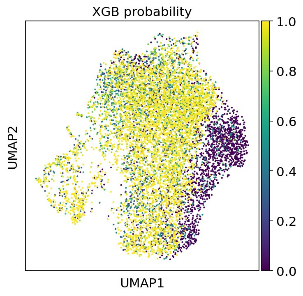

**Supplementary Figure 11:** **UMAP visualization of the predicted male probabilities the cells assigned by different classifiers in the chimeric mouse diaphragm experimental dataset.** From left to right: Random Forest, SVM, Logistic Regression, and XGBoost classifiers. The color gradient ranges from purple (low probability) to yellow (high probability).

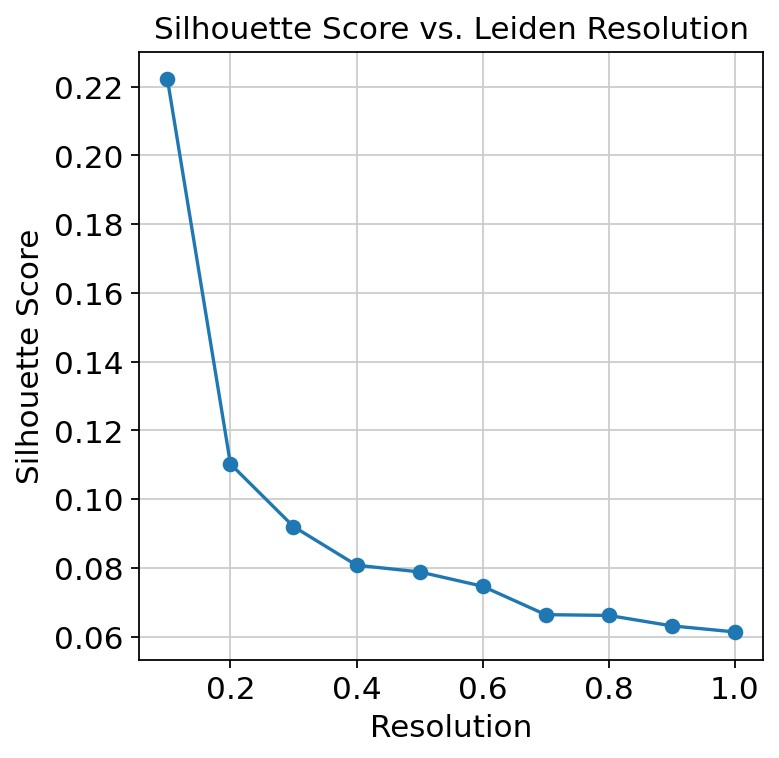

**Supplementary Figure 12:** **Evaluation of Leiden clustering resolution on the chimeric mouse diaphragm experimental dataset using Silhouette score.** The Silhouette score was computed across a range of Leiden resolution parameters to quantitatively assess clustering quality, with higher scores indicating more compact and well-separated clusters. We observed a rapid decline in Silhouette score at lower resolutions, followed by a gradual decrease. To balance between maximizing cluster separation and capturing biological heterogeneity, we selected a resolution of 0.6, just before a more pronounced score drop. This approach ensures the stability and interpretability of downstream analyses while avoiding over-partitioning of cell populations.

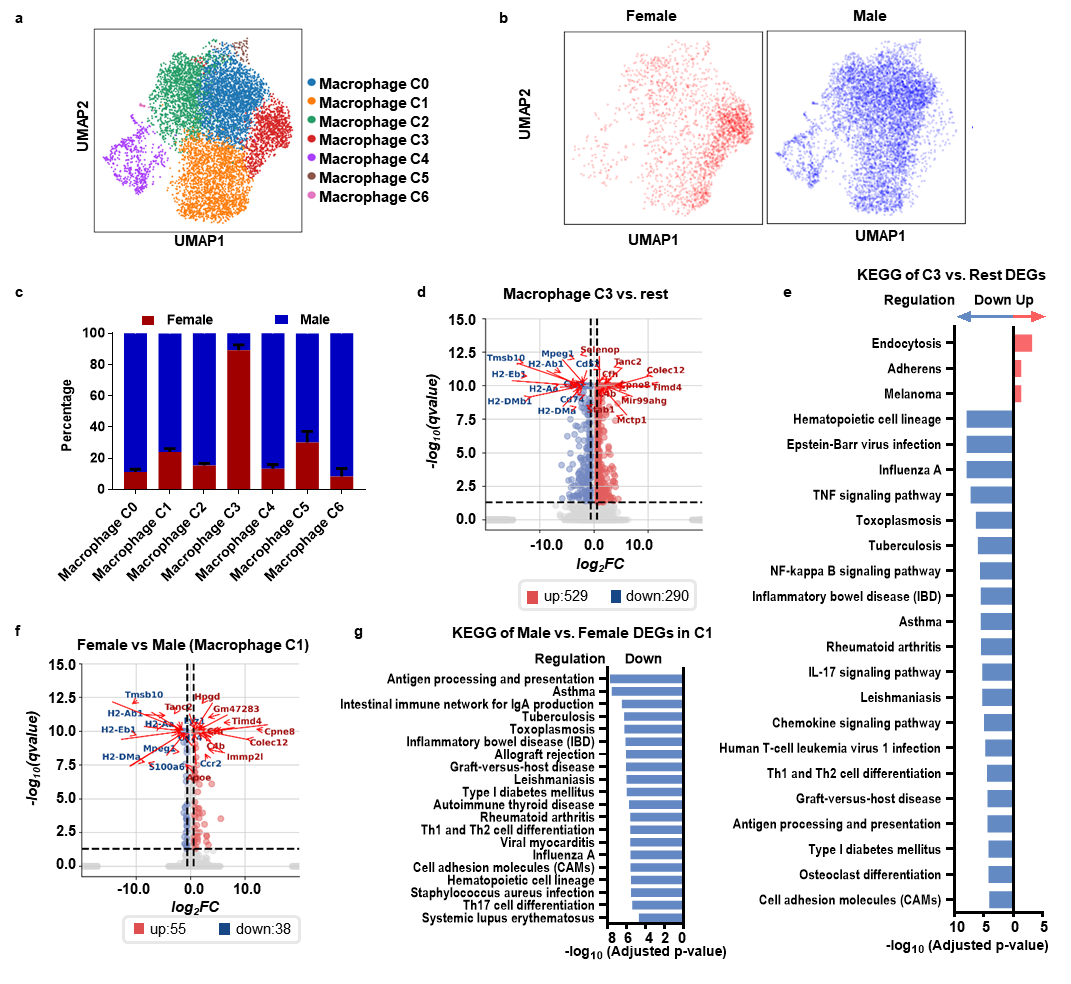

**Supplementary Figure 13:** **Chimeric mouse diaphragm experimental dataset analysis results at resolution = 0.4. a,** UMAP visualization showing macrophage clusters (C0–C6). **b,** UMAP plots colored by sex (Female and Male). **c,** Proportion of female and male cells in each macrophage cluster. **d,** Volcano plot of differentially expressed genes in Macrophage C3 versus the rest of the clusters (upregulated genes in red, downregulated in blue). **e,** Pathway enrichment analysis of DEGs in Macrophage C3, showing upregulated (red) and downregulated (blue) pathways. **f,** Volcano plot comparing female and male cells within Macrophage C1. **g,** Pathway enrichment analysis for sex-biased DEGs in Macrophage C1, showing downregulated pathways in female cells.

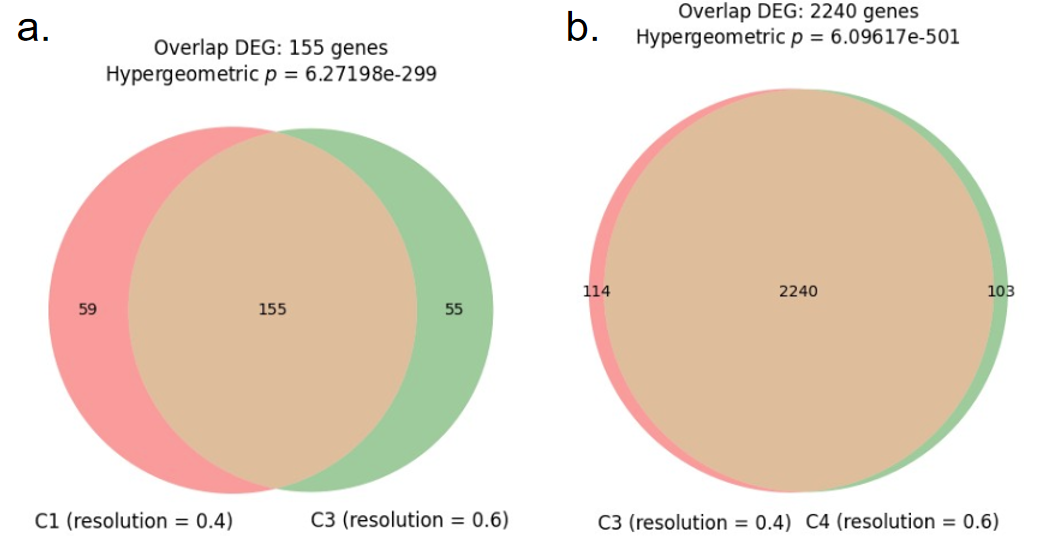

**Supplementary Figure 14:** **Comparison of the DEG analysis following cluster resolution 0.6 and 0.4 in the chimeric mouse diaphragm experimental dataset.** DEGs between corresponding clusters in two versions are highly similar in the two versions. a, Overlap of DEGs from the female versus male comparison within cluster C3 at resolution 0.4 and cluster C4 at resolution 0.6. The overlap consisted of 155 genes between two DEG sets (hypergeometric test: *p* = 6.27 × 10⁻²⁹⁹). b, Overlap of DEGs from cluster-versus-rest comparisons for cluster C1 at resolution 0.4 and cluster C3 at resolution 0.6. The overlap included 2,240 shared DEGs (hypergeometric test: *p* = 6.10 × 10⁻⁵⁰¹).

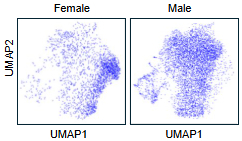

**Supplementary Figure 15: UMAP visualizations of the male and female cells in the chimeric mouse diaphragm experimental dataset with monotonic color.** UMAP visualizations showing the spatial distribution of recipient- and donor-derived macrophages from the chimeric mouse diaphragm experimental datasets, using the same color to facilitate distribution comparison.

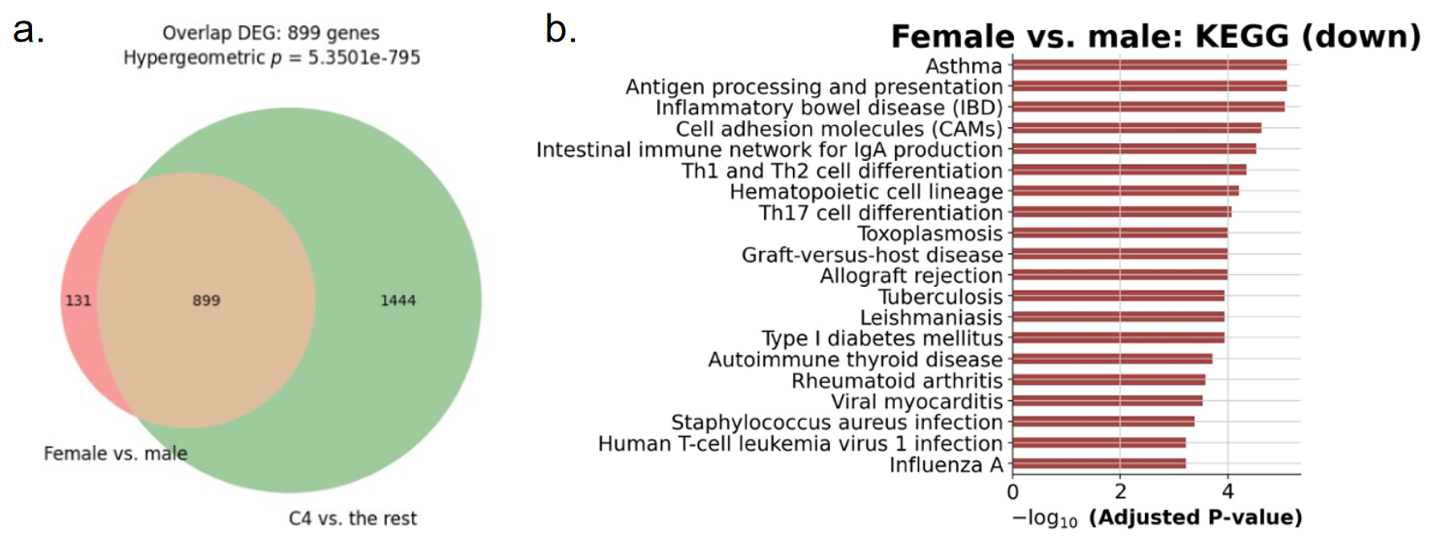

**Supplementary Figure 16: Sex-specific DEGs significantly overlap with C4 cluster DEGs and enrich immune-related pathways in the chimeric mouse diaphragm dataset. a,** Venn diagram illustrating the overlap between DEGs from the C4 cluster vs. the rest comparison and global female vs. male DEGs, revealing 899 shared genes (hypergeometric *p*-value = 5.35 × 10⁻⁷⁹⁵). **b,** KEGG pathway enrichment analysis of genes downregulated in females relative to males. Several immune-related pathways are significantly enriched, including antigen processing and presentation, inflammatory bowel disease (IBD), cell adhesion molecules (CAMs), the intestinal immune network for IgA production, and Th1 and Th2 cell differentiation. Notably, these pathways overlap with those enriched in the C4 vs. rest analysis shown in Figure 6 (panel g), further supporting shared immune modulation and antigen presentation mechanisms that underlie sex differences across macrophage subtypes and analysis resolutions.

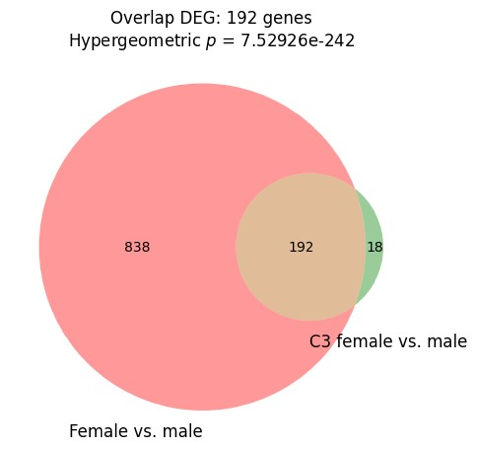

**Supplementary Figure 17:** **Overlap analysis of differentially expressed genes (DEGs) in the chimeric mouse diaphragm experimental dataset**. Venn diagram illustrating the overlap between DEGs from C3 female vs. male analysis and global female vs. male DEGs, revealing 192 shared genes (hypergeometric *p*-value = 7.53 × 10⁻²⁴²). (Supplementary figure 16, panel a) Venn diagram showing the overlap between DEGs identified in the comparison of C4 cluster vs. the rest and DEGs from female vs. male analysis. A total of 899 overlapping genes were observed, with a hypergeometric *p*-value of 5.35 × 10⁻⁷⁹⁵, indicating a significant overlap.

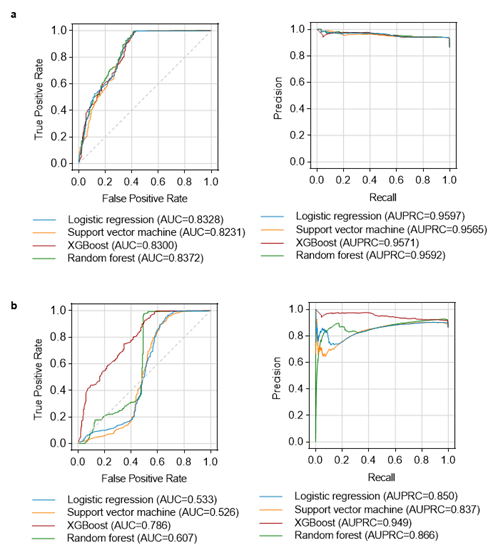

**Supplementary Figure 18: Batch effect correction can reduce biological signal, leading to decreased prediction performance.** To evaluate the impact of batch effect correction on biological signal retention, we applied Scanorama to correct batch effects in the mouse adrenal gland dataset. This corrected dataset was then used to train machine learning models based on 14 sex-linked markers, which were subsequently tested on the chimeric mouse diaphragm validation dataset. The performance is then compared with the models trained on the same dataset without batch effect correction. Compared to the uncorrected version (panel **a**), models trained on the batch-corrected data show markedly reduced prediction performance (panel **b**), suggesting that batch effect correction removed biologically relevant variation critical for accurate sex prediction.
